# RIF1 regulates replication origin activity and early replication timing in B cells

**DOI:** 10.1101/2023.03.31.535086

**Authors:** Daniel Malzl, Mihaela Peycheva, Ali Rahjouei, Stefano Gnan, Kyle N. Klein, Mariia Nazarova, Ursula E. Schoeberl, David M. Gilbert, Sara C. B. Buonomo, Michela Di Virgilio, Tobias Neumann, Rushad Pavri

## Abstract

The mammalian DNA replication timing (RT) program is crucial for the proper functioning and integrity of the genome. The best-known mechanism for controlling RT is the suppression of late origins of replication in heterochromatin by RIF1. Here, we report that in antigen-activated B lymphocytes, RIF1 binds predominantly to early-replicating active chromatin, regulates early origin firing and promotes early replication. RIF1 has a minor role in gene expression and genome organization in B cells. Furthermore, we find that RIF1 functions in a complementary and non-epistatic manner with minichromosome maintenance (MCM) proteins to establish early RT signatures genome-wide and, specifically, to ensure the early replication of highly transcribed genes. These findings reveal new layers of regulation within the B cell RT program, driven by the coordinated activity of RIF1 and MCM proteins.

## INTRODUCTION

The faithful and timely replication of the genome is essential for inheriting genetic information and avoiding chromosomal abnormalities. To ensure this, large metazoan genomes initiate replication from several discrete loci, termed origins of replication. Origins are not activated simultaneously across the genome, but rather, in an asynchronous manner referred to as the DNA replication timing (RT) program^1^. A hallmark of the RT program is that genomic A compartments enriched in transcriptionally active genes generally replicate earlier in S phase whereas B compartments harboring silent heterochromatin typically replicate later in S phase^2, 3^. Deregulation of RT has been correlated with defects in chromosome condensation, sister chromatid cohesion, gene expression and genome instability ^4–6^. Altered RT can disrupt the distribution of active and repressive epigenetic marks causing major alterations in genome architecture^7^. In genetic diseases and cancer, defects in RT have been correlated with deleterious chromosomal translocations^8–12^. Recently, RT has been directly implicated in the biogenesis of oncogenic translocations found in B cell lymphomas and other leukemias^13^. In addition, late replication has been consistently associated with higher rates of mutation across species ^11, 14–17^ and stress-induced delays in replication are a hallmark of common fragile sites in long, transcribed genes^18–20^. Yet, despite its key role in maintaining cellular physiology and genome integrity, our understanding of the mechanisms regulating RT remains incomplete.

Origins are specified in the late G2/M and early G1 phases of the cell cycle via the loading of the origin recognition complex (ORC) and associated factors^21–25^. The subsequent recruitment of the hexameric ring-shaped MCM complex helicase (MCM2-7) licenses these sites for activation^22^, resulting in the formation of the pre-replication complex (pre-RC)^21–25^. In S phase, a subset of licensed origins is activated by a set of proteins collectively called replication firing factors, resulting in the initiation of replication^21–24^. A subset of licensed, MCM-bound origins can be activated by firing factors at any given point in S phase^26^. This has led to the idea that firing factors must be recycled to activate subsequent sets of origins later in S phase, thus ensuring timely completion of replication. In support, overexpression of firing factors has been shown to advance the RT of late-replicating chromatin^27–29^. These studies imply that the time of recruitment of the firing factors to a licensed origin determines its RT ^30^. In essence, the probability of origin activation in S phase determines the RT of a region, meaning that early RT domains harbor origins that typically fire earlier in S phase whereas late RT domains contain origins that tend to fire later in S phase.

The best-studied mechanism of RT control is the suppression of late origin firing by the multifunctional protein, RIF1^31–33^. RIF1 associates with late-replicating chromatin in large, mega-base domains called RIF1-associated domains (RADs) and represses origin firing via protein phosphatase 1 (PP1)-mediated dephosphorylation of MCM4^34–36^. Ablation of RIF1 (*Rif1*^-/-^) in human and mouse embryonic stem cells (hESCs and mESCs) and in primary mouse embryonic fibroblasts ^31^ resulted in a genome-wide loss of early and late RT domain distinction associated with alterations in epigenetic marks and genome compartmentalization^7, 35, 36^. However, in some *Rif1*^-/-^ cell lines, this RT phenotype was considerably weaker ^7^, suggesting the existence of additional modes of RT control.

We recently showed that in MCM-depleted CH12 cells, a murine B cell line, the RT program was globally deregulated without major changes in transcription or genome architecture^13^. Since RIF1 is the only other factor whose loss could lead to such a major phenotype^7, 31, 35^, we investigated the role of RIF1 in in the B cell RT program. We report the surprising findings that RIF1 is predominantly bound to active chromatin in B cells, regulates early origin activity in these regions and promotes their early replication. In addition, we find that RIF1 acts in a complementary and non-epistatic manner with MCM complexes to drive early replication, especially of highly transcribed regions. In sum, our study reveals an additional regulatory layer within the global RT program and a new role for RIF1 in promoting early replication.

## RESULTS

### RIF1 regulates early replication in B cells

To measure RT, we performed Repli-seq from early (E) and late (L) S phase fractions in normal and *Rif1*^-/-^ CH12 B cells^37, 38^ (Fig. S1A). MCM complexes were depleted by infection with lentiviruses expressing short hairpin RNAs (shRNAs) targeting Mcm6 (shMcm6), and an shRNA against LacZ (shLacZ) served as a control for infection and shRNA expression, as described previously^13, 39^ (Fig. S1B). RT was calculated as the (log2) E/L ratio^40^ (Fig. 1A). RT values showed the expected bimodal distribution in shLacZ cells reflecting distinct early (E; positive RT) and late (L; negative RT) domains (Fig. 1A), which was also reflected in the genomic RT profiles (Fig. 1B). In shMcm6 cells, this distinction was globally weakened with all early and late RT values approaching zero (Fig. 1A-B). In

**Figure 1:**
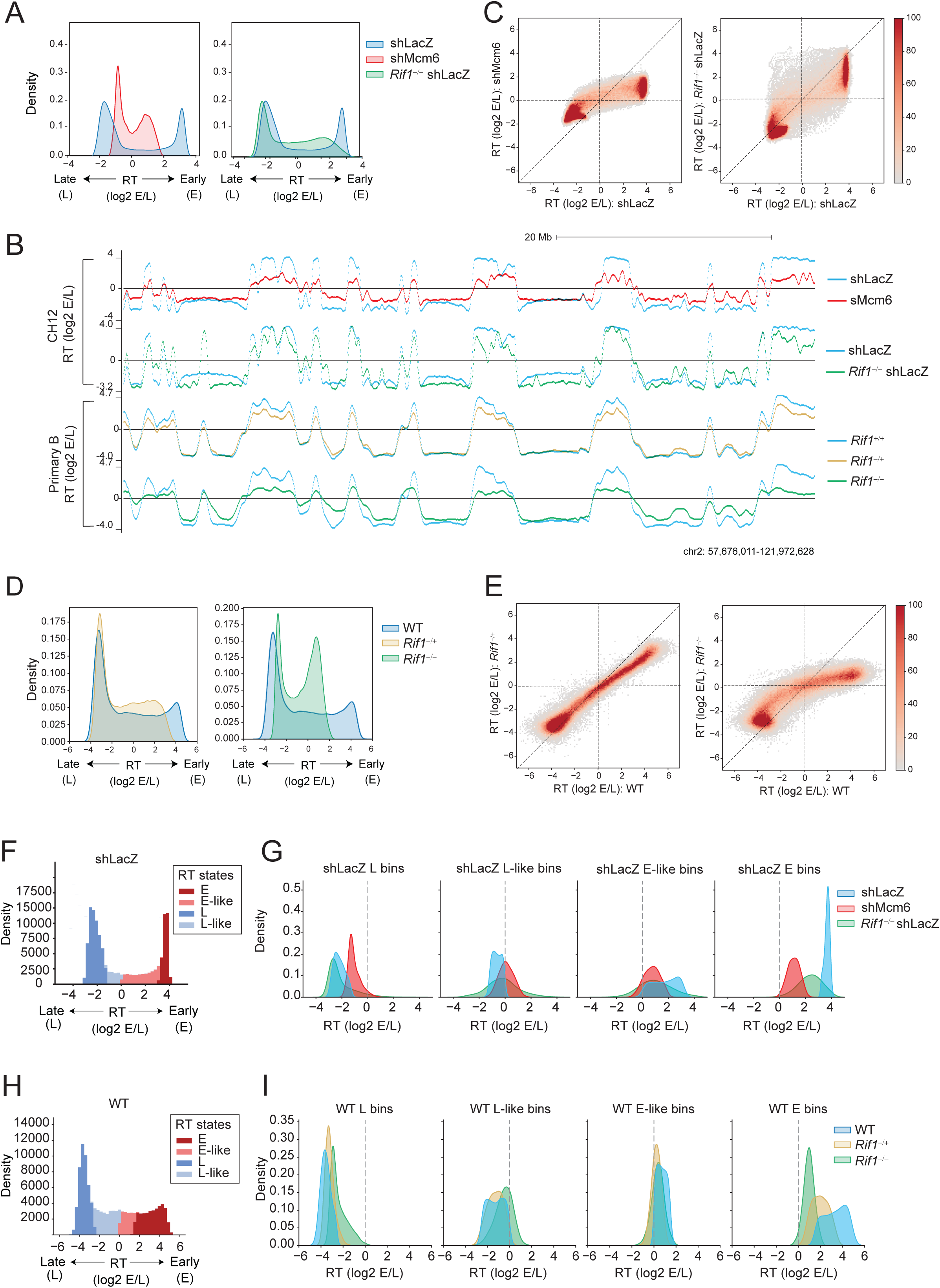
RIF1 regulates early RT in activated B cells. A. RT histograms in 5 kb genomic bins from shLacZ (blue), shMcm6 (red) and *Rif1*^-/-^ shLacZ (green) CH12 cells. RT is calculated as the ratio of the read densities of early (E) and late (L) Repli-seq fractions (log2 E/L). B. UCSC genome browser view of RT for the indicated conditions in CH12 cells (top panel) and primary, activated splenic B cells (bottom panel). Positive and negative values correspond to early and late-replicating regions. C. Comparison of RT values in 20 kb genomic bins between shLacZ and shMcm6 CH12 cells (left) and shLacZ and *Rif1*^-/-^ shLacZ cells (right). D. RT histograms as in (A) from WT (blue), *Rif1*^-/+^ (ochre) and *Rif1*^-/-^ (green) primary, activated splenic B cells. E. Comparison of RT values in 20 kb genomic bins between WT, *Rif1*^-/+^ and *Rif1*^-/-^ in primary B cells. F. Classification of shLacZ RT bins into four states using a Hidden Markov Model (HMM). Three states E, M (mid) and L were called, and the middle state was split into into E-like and L-like at the log2 (E/L) = 0 boundary. G. RT histograms showing how the bins in each shLacZ RT state (blue) called in (D) change in shMcm6 (red) and *Rif1*^-/-^ shLacZ (green) CH12 cells. H. HMM analysis as in F but using WT RT values from primary B cells to call RT states. I. RT histograms showing how bins in each WT RT state (blue) change in *Rif1*^-/+^ (ochre) and *Rif1*^-/-^ (green) primary B cells.

*Rif1*^-/-^ shLacZ cells, many early-replicating domains underwent a delay in replication (Fig. 1B). In comparison, late RT domains showed a mixed phenotype in *Rif1*^-/-^ shLacZ cells with some, typically smaller, domains showing advanced replication signatures, and other, typically larger, domains undergoing delayed replication relative to shLacZ cells (Fig. 1A-B). Direct comparison of RT in 20 kb genomic bins between the different conditions showed that most early- and late-replicating bins were strongly shifted towards zero in shMcm6 cells indicative of a deregulation of the RT program (Fig.1C). In *Rif1*^-/-^ shLacZ cells, most early-replicating regions underwent delayed replication to varying degrees (note the red stripe in Fig. 1C), albeit to a lesser extent than in shMcm6 cells. Although late-replicating bins were mildly affected in *Rif1*^-/-^ shLacZ cells, many of these bins underwent slightly delayed RT (Fig. 1C). The latter observation is reminiscent of reports in other *Rif1*^-/-^ cell lines where the RT of many late-replicating domains that were not associated with RIF1 was unaffected by the loss of RIF1 ^7^.

We also performed Repli-seq in a more physiological system, namely, primary activated splenic B cells from *Rif1*^+/+^, RIF1 heterozygous (*Rif1^-/+^*) and *Rif1*^-/-^ mice^41^ (Fig. S1C). WT primary B cells had relatively fewer early replicating domains and more mid-replicating domains than shLacZ CH12 cells, resulting in different RT density profiles (Fig. 1D, compare with Fig. 1A). Strikingly, we observed a nearly exclusive effect on early-replicating genomic bins in heterozygous *Rif1*^-/+^ cells indicating that reduced levels of RIF1 protein was sufficient to delay replication of early-replicating regions without majorly affecting late replicating ones (Fig. 1 D-E). This phenotype was exacerbated in homozygous *Rif1^-/-^* cells as seen by the further delay in replication of early RT domains, which was accompanied by the earlier replication of several late RT domains albeit to varying degrees (Fig. 1 B and D-E). These results, especially from heterozygous cells, highlight a new role for RIF1 in regulating early replication in B cells.

To systematically measure changes in RT upon loss of RIF1, we generated three RT states, early (E), middle (M) and late (L), in shLacZ CH12 cells using a Hidden Markov Model (HMM) to segment the genome into 20 kb bins based on their RT value (Fig. 1F). We split the M state at the zero RT value into early-like (E-like) having positive RT values and late-like (L-like) having negative RT values, which resulted in four RT classes (Fig. 1F). To allow direct comparison of RT changes between conditions, we created density plots for each of the four RT states in shLacZ cells displaying the RT of all other experimental conditions in those genomic bins. The results showed that E and E-like bins in shLacZ replicated later in *Rif1*^-/-^ cells, but, importantly, that this was not accompanied by commensurate advances in the RT of L bins suggesting that loss of RIF1 does not globally deregulate RT (Fig. 1G). Indeed, most L bins retained their RT values with only some bins showing advanced RT and some others showing further delays in RT relative to shLacZ cells (Fig. 1G). In contrast, in shMcm6 cells, all shLacZ RT bins shifted towards zero RT values implying that these cells have undergone a global deregulation of the RT program (Fig. 1G). The same HMM-based analysis in primary, splenic B cells revealed that in *Rif1^-/+^* heterozygotes, there was a strong delay in the RT of E bins associated with a much weaker advance in the RT of L bins and virtually no changes in L-like bins, which further demonstrates that RIF1 primarily regulates early replication in B cells (Fig. 1H-I). These changes were exacerbated in *Rif1*^-/-^ B cells as seen by the further delay in the RT of E bins which as accompanied by the advanced RT of L bins relative to *Rif1^-/+^* cells (Fig. 1H-I).

We conclude that RIF1 functions as modulator of early replication in B cells rather than as a global regulator of RT.

### RIF1 plays a minor role in gene expression and genome compartmentalization in B cells

To determine the role of RIF1 in gene expression, we performed RNA-seq from two independent clones (clones 1 and 2) of *Rif1*^-/-^ CH12 cells and identified differentially regulated genes with the DESeq2 software ^42^ (Fig. S2A-B). Only 22 downregulated and 3 upregulated genes were common between the two clones. A gene ontology (GO) analysis of the common downregulated genes did not reveal enrichments in pathways related to DNA replication, cell division, cell proliferation or genome organization (Fig. S2C), and there were no pathways enriched in the common upregulated genes.

RIF1 has been implicated in regulating genome compartmentalization, in part, through deregulation of RT ^7^. However, in *Rif1^-/-^* shLacZ CH12 cells, Hi-C analysis revealed no gross changes in compartment profiles based on evaluation of Hi-C heatmaps where compartments manifest as the checkerboard patterns off the diagonal (Fig. 2A). We quantified the compartment signals based on the first principal component (PC1) of the Hi-C contact matrix where positive PC1 values denote active, early-replicating A compartments and negative PC1 values denote silent, late-replicating B compartments (Fig. 2B-C)^43^. Comparative PC1 analysis showed that although the majority of the bins in *Rif1*^-/-^ shLacZ cells underwent minor changes in their PC1 values relative to shLacZ cells, a few bins did shift substantially, including PC1 sign flips in both directions, which gave a more dispersed appearance to the distribution relative to control cells (Fig. 2B, right). Visualization of PC1 profiles in various genomic regions confirmed that small changes in PC1 occurred in many locations in *Rif1*^-/-^ shLacZ cells, but importantly, that these did not always correlate with the changes in RT (Fig. 2C). However, the magnitude of PC1 changes and the frequency of compartment switching (PC1 sign flips) is considerably milder than that reported in other cell types where loss of RIF1 led to substantial compartment switching^7, 35^. In comparison, PC1 values in shMcm6 cells were comparable to those in shLacZ cells (Fig. 2B, left and Fig. 2C)^13^.

**Figure 2:**
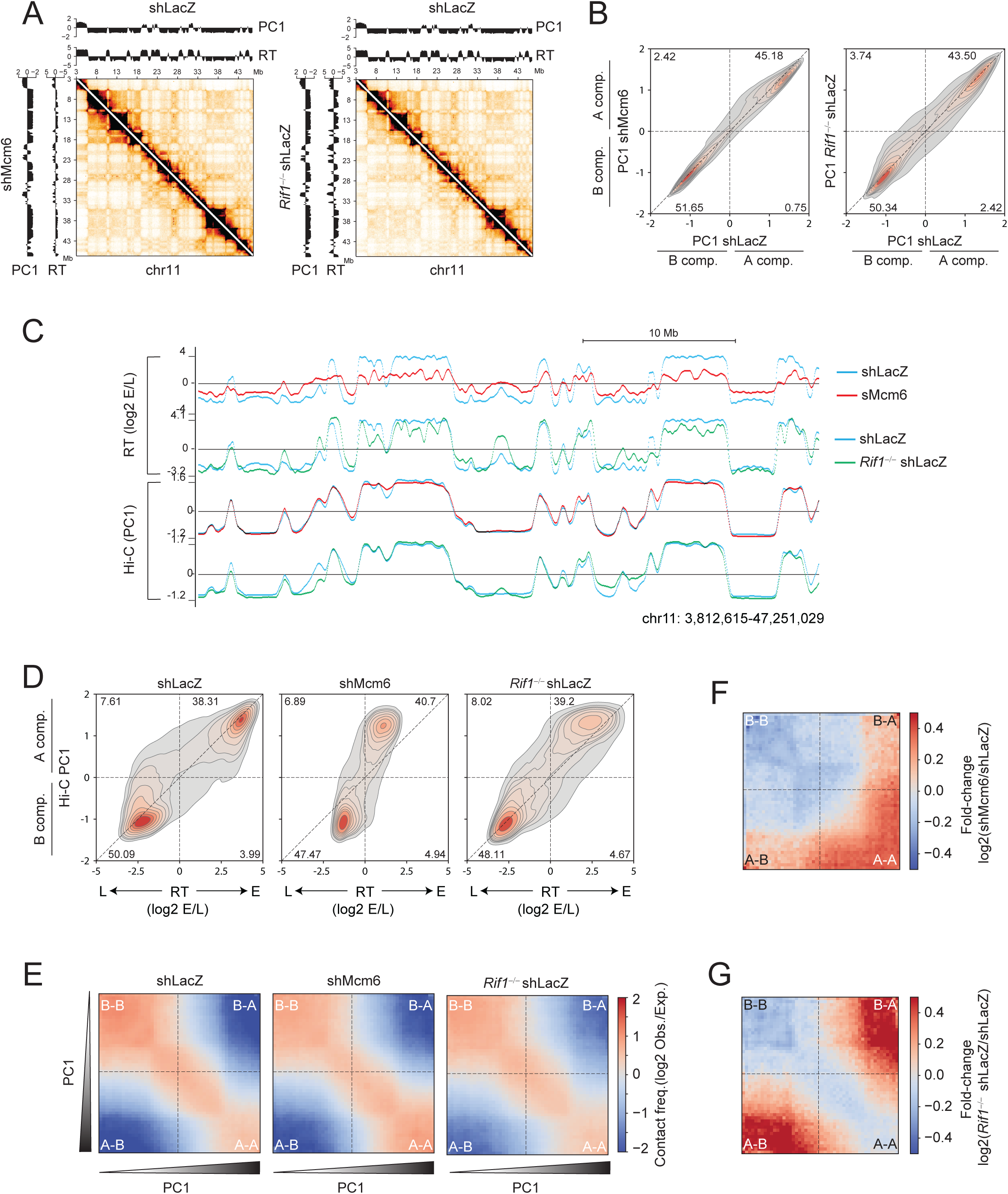
RIF1 has a minor role in B cell genome architecture. A. <u>Left</u>: Hi-C contact matrix of chromosome 11 showing shLacZ contacts (above the diagonal) and shMcm6 contacts (below the diagonal). The PC1 compartment signals (Hi-C eigenvector eigenvalues) and RT (log2 E/L) tracks are shown above (for shLacZ) and on the left (for shMcm6) for each matrix. <u>Right</u>: Same as before but comparing shLacZ and *Rif1*^-/-^ shLacZ cells. B. Density-contour plot of PC1 compartment signals per 20 kb genomic bin. The left plot compares the PC1 signals in shLacZ and shMcm6 cells, and the right plot compares PC1 values between shLacZ and *Rif1*^-/-^ shLacZ cells. The numbers within the plots are the percentage of bins in that quadrant. C. A representative UCSC genome browser view comparing Hi-C PC1 profiles with RT (log2 E/L) profiles in shLacZ, shMcm6 and *Rif1*^-/-^ shLacZ cells. D. PC1 versus RT density-contour plots in 20 kb genomic bins to compare changes in compartmental identities (PC1) with RT in shLacZ, shMcm6 and *Rif1*^-/-^ shLacZ CH12 cells. E. Saddle plot from shLacZ, shMcm6 and *Rif1*^-/-^ shLacZ CH12 cells showing long-range (> 2Mb) intra- chromosomal contact enrichments between bins of varying compartment signal strength (PC1). The values were computed from 20 kb KR-normalized contact matrices. F-G Fold-change saddle plots highlighting the changes in compartmentalization in shMcm6 and *Rif1*^-/-^ shLacZ cells relative to shLacZ cells.

We also generated PC1 versus RT density contour plots which allowed us to simultaneously compare the changes in compartmentalization and RT within the same set of genomic 20 kb bins. The density contour distribution profile in *Rif1^-/-^* shLacZ cells showed a marked shift of early RT bins towards delayed RT but without major changes in their PC1 values (Fig. 2D). This implies that the roles of RIF1 in RT and genome architecture are separable in B cells, as they are in other cells^7, 35^. However, *Rif1*^-/-^ shLacZ cells showed a gain of A-B inter-compartmental interactions with a corresponding decrease in A-A and B-B contacts (Fig. 2E-F). This effect was clearly manifested in fold-change saddle plots where contacts between regions of highest PC1 (early-replicating) and lowest PC1 (late-replicating) were increased in *Rif1*^-/-^ shLacZ cells (Fig. 2F). Thus, RIF1 is important for maintaining normal compartmentalization in B cells by preventing the mixing of A and B compartments. Importantly, this compartmentalization phenotype was observed in mESCs and hESCs where, in contrast to B cells, loss of RIF1 caused a severe deregulation of the RT program^7, 35^. This suggests that the changes in compartmentalization in *Rif1*^-/-^ cells are conserved between diverse cell lineages, but that these are unrelated to the changes in RT. Moreover, in shMcm6 cells, where RT is globally disrupted to a similar degree as in *Rif1*^-/-^ mESCs and hESCs ^7, 13, 35^, we observed relatively minor changes in compartmentalization, as seen by the mild increase in A-A contacts and a similar decrease in B-B contacts (Fig. 2E-F).

In sum, the uncoupling of RT from genome organization in B cells and other cells ^7, 13, 35^ leads us to conclude that although RIF1 contributes to the normal spatial separation of A and B compartments in B cells, it’s role in regulating early RT is unlikely to be directly linked to these structural changes.

### RIF1 is predominantly located in early-replicating transcribed chromatin in B cells

We next investigated the genomic occupancy of RIF1 in B cells. We performed chromatin immunoprecipitation (ChIP) from primary, mature splenic B cells derived from the *Rif1*^FH/FH^ mouse line wherein RIF1 is endogenously tagged at its C terminus with the Flag and Hemagglutinin (HA) epitopes^31, 36^. RIF1 chromatin occupancy was determined via ChIP sequencing (ChIP-seq) with an anti-HA antibody in *Rif1*^FH/FH^ and *Rif1*^+/+^ cells (Fig. S3A).

Visual analysis on the genome browser revealed RIF1 binding predominantly in early replicating domains in B cells (Fig. 3A). Comparison with ChIP-seq profiles from *Rif1*^FH/FH^ mouse embryonic fibroblasts (MEFs)^36^ showed that while RIF1 localized to broad domains in late-replicating regions in MEFs, in B cells, enrichments were mostly in early-replicating domains (Fig. 3B). We next identified peaks of RIF1 occupancy ^44^ as well as RADs^45^ in both B cells and MEFs. Peaks in both cell types were enriched in early-replicating regions (Fig. 3C-D). However, although RADs in MEFs were mostly late-replicating, as described^36^, in B cells, they were largely early-replicating (Fig. 3C-D).

**Figure 3:**
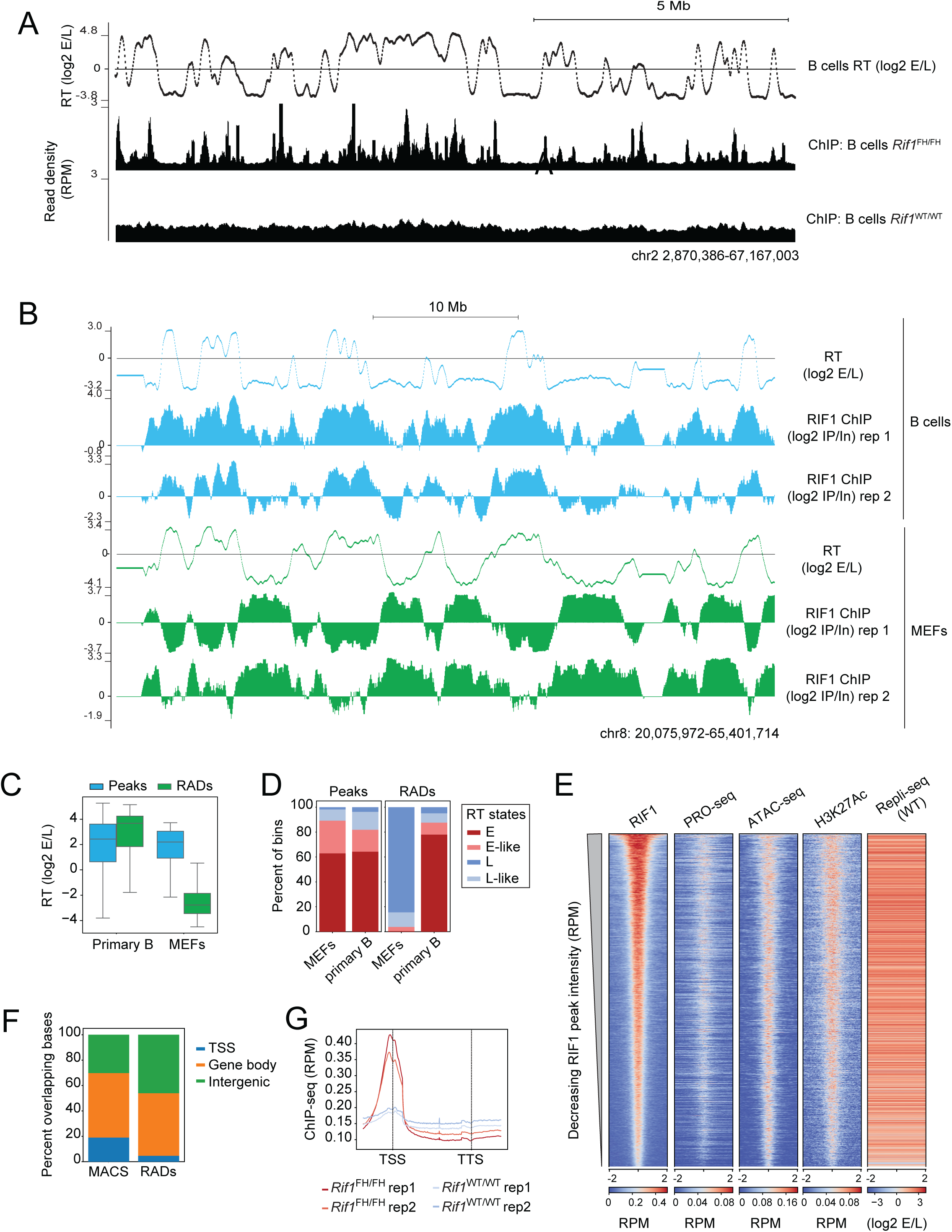
RIF1 localizes predominantly to active chromatin in activated B cells. A. A representative UCSC genome browser view of RIF1 occupancy in *Rif1*^FH/FH^ primary, activated splenic B cells. ChIP-seq was performed with an anti-HA antibody and *Rif1*^WT/WT^ cells were used as a negative control. The RT track from WT, primary B cells provides a reference for early and late- replicating domains. B. UCSC browser snapshot comparing ChIP enrichments of RIF1 in primary B and MEFs from *Rif1*^FH/FH^ mice. To allow direct comparison, ChIP signal was normalized to input signal and the ratio (ChIP/Input) tracks are shown. RT tracks are from WT cells. C. Box plots comparing RT values of RIF1 peaks and RADs in MEFs and primary B cells. 16,043 peaks and 289 RADs were identified in primary B cells and 862 peaks and 332 RADs were called in MEFs. D. Peaks and RADs were called as in C above and classified into the four RT states described in Fig. 1D. E. Heatmap analysis of the chromatin locale in a 4 kb window surrounding RIF1 peaks called in *Rif1*^FH/FH^ cells. The heatmap is centered on the RIF1 peak summit and ordered by decreasing RIF1 read density (reads per million, RPM). All other genomic tracks shown (PRO-seq, ATAC-seq, H3K27Ac and Repli-seq) are from WT primary, activated splenic B cells. F. Distribution of RIF1 peaks and RADs at TSSs, gene bodies and intergenic regions. G. Metagene analysis showing average RIF1 occupancy patterns at genes in *Rif1*^FH/FH^ and *Rif1*^WT/WT^ cells from two replicates each (rep1 and rep2). The TSS and TTS are the transcription start site and transcription termination site.

We next determined the overlap of RIF1 peaks with nascent transcription measured by precision run-on sequencing (PRO-seq)^46^, chromatin accessibility, measured by assay for transposase-accessible chromatin (ATAC-seq)^47^, and ChIP-seq of histone H3 acetylated at lysine 27 (H3K27Ac), a mark of active transcription start sites (TSSs) and enhancers. The resulting heatmaps, centered on the RIF1 peak summits, revealed an enrichment of RIF1 in transcribed, H3K27ac-rich accessible chromatin (Fig. 3E). In agreement, ∼70% of RIF1 peaks and ∼50% of RADs overlapped TSSs or gene bodies (Fig. 3F), and a metagene analysis showed that RIF1 was relatively enriched at TSSs (Fig. 3G).

We conclude that RIF1 predominantly occupies early-replicating, transcribed chromatin in B cells, suggesting that RIF1 promotes their early replication via direct association.

### RIF1 regulates the activity of a subset of origins of replication in active chromatin

The localization of RIF1 in active chromatin coupled with the delayed replication of these domains in *Rif1^-/-^* cells led us to investigate whether RIF1 played a role in promoting origin firing in activated B cells. To address this, we measured origin activity with short nascent strand sequencing (SNS-seq) in WT and *Rif1^-/-^* CH12 cells (Fig. S3B). SNS-seq maps the location and relative usage of origins (termed replication initiation sites; ISs) in a population of cells by quantifying the levels of nascent leading strands^48, 49^. As described in our previous work ^13^, ISs and initiation zones were defined based on peak-calling and peak-clustering, respectively, and only ISs within initiation zones were used for analysis.

We generated heatmaps of IS read densities in a 4 kb window centered at the IS peak summit and ordered by the IS density in WT cells (Fig. 4B). To visualize how IS density correlated with various chromatin features in the genomic neighborhood, we also generated heatmaps showing the densities of DNase hypersensitive sites (DHSs) which mark accessible chromatin at active promoters and enhancers, H3K36me3, which is enriched in the bodies of transcriptionally active genes, and the repressive mark, H3K9me3 (Fig. 4B). In WT cells, the most active ISs were embedded in active chromatin and gene bodies whereas the least active ISs were associated with H3K9me3 (Fig. 4B). In *Rif1^-/-^* cells, origin activity was repressed in active chromatin but increased in H3K9me3 domains (Fig. 4B). This phenotype is reminiscent of what we previously reported in shMcm6 B cells ^13^. Importantly, however, only ∼5% of ISs are deregulated (1.5-fold) upon loss of RIF1 compared to ∼38% in shMcm6 cells (Fig. S3C), which correlates with the magnitude of their respective RT phenotypes (Fig. 1).

**Figure 4:**
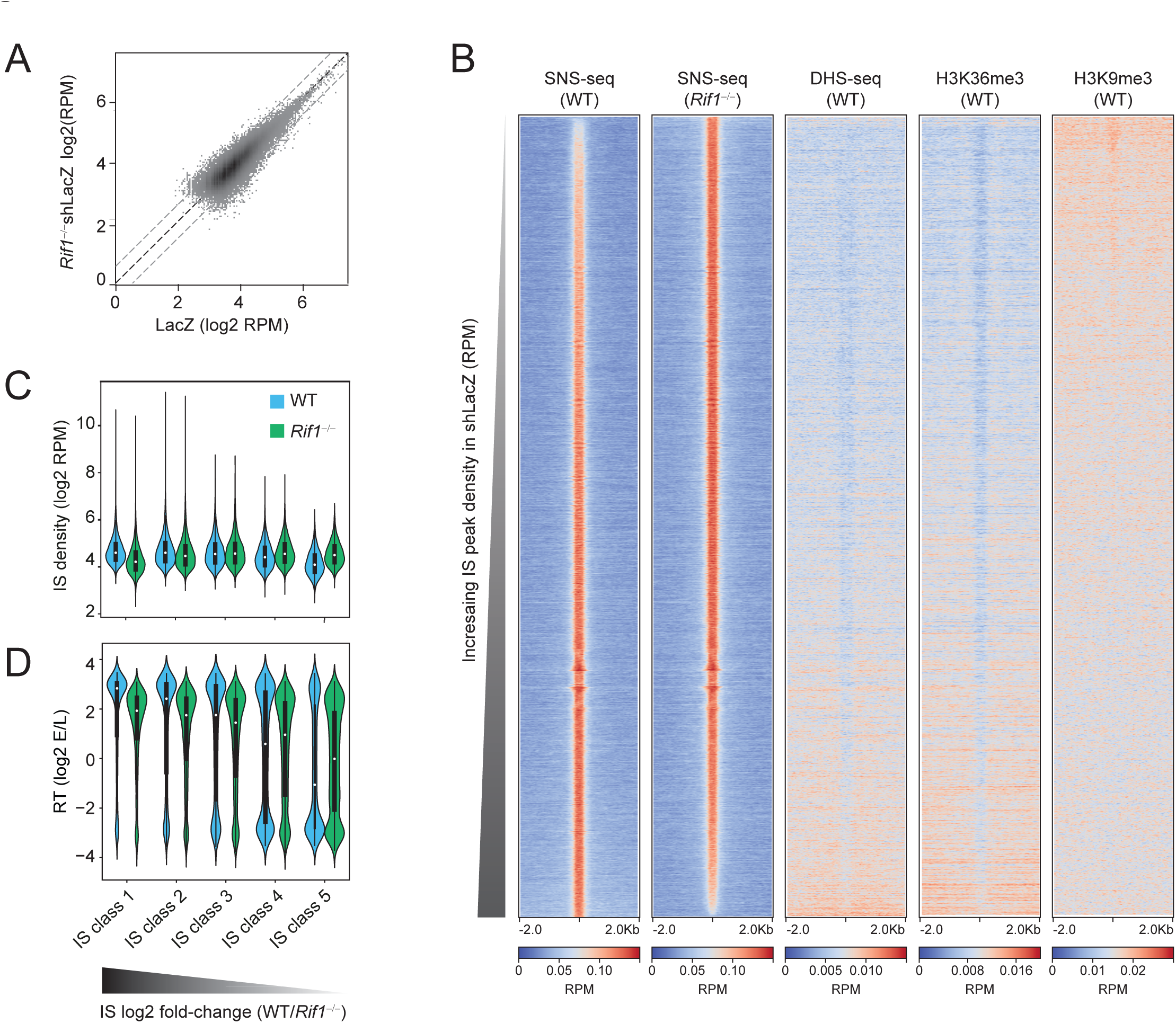
RIF1 regulates the activity of early origins of replication in B cells. A. Scatter plot of IS density in log2 RPM (reads per million) in WT and *Rif1*^-/-^ cells identified from SNS-seq. B. Heatmaps spanning a 4 kb region centered on the IS peak summit were generated to visualize the profiles of SNS-seq, DNase hypersensitivity (DHS)-seq and histone modifications (H3K36me3 and H3K9me3) around ISs. All heatmaps display read densities as RPM. Within each heatmap, the ordering is based on decreasing SNS-seq fold-change (*Rif1*^-/-^/WT). C. ISs were split into five equal classes (quintiles) based on their fold-change (log2 *Rif1*^-/-^/WT) such that class 1 contained the most downregulated ISs and class 5 contained the most upregulated ISs, respectively, in *Rif1*^-/-^ cells. The violin plots show the IS density within each class in WT and *Rif1*^-/-^ cells. D. ISs were classified as in C above. The violin plots show the RT values within each IS class.

To quantify the changes in origin efficiency, we calculated fold-changes of ISs (log2 WT/ *Rif1^-/-^*), ranked them from highest to lowest and created five equal classes (quintiles) such that class 1 contained the most downregulated ISs in *Rif1^-/-^* cells and class 5 contained the most upregulated ISs in *Rif1^-/-^*cells. We next generated violin plots displaying the density of ISs (Fig. 4C) or RT values (Fig. 4D) in WT and *Rif1^-/-^* cells within each IS class. The results showed that in WT cells, class 1 ISs were the most active (Fig. 4C) and were mostly early-replicating (Fig. 4D) whereas class 5 ISs were the least active with the majority being late-replicating (Fig. 4C-D). Importantly, class 5 IS densities in *Rif1^-/-^* cells were comparable to class 1 IS densities in WT cells, suggesting that the upregulated origins in *Rif1^-/-^* cells (class 5), which are normally the weakest, fire at similar efficiencies as the most active origins in WT cells (class 1) (Fig. 4C).

We conclude that RIF1 is required for the optimal activity of a subset of early replicating origins in B cells.

### RIF1 and MCM complexes act in a complementary manner to regulate the B cell RT program

The differing RT phenotypes in *Rif1^-/-^* and shMcm6 cells led us to hypothesize that they may function in a non-epistatic manner to drive early replication in B cells. To address this, we infected *Rif1*^-/-^ cells with lentiviruses expressing shMcm6 (*Rif1^-/-^* shMcm6) or shLacZ (*Rif1^-/-^* shLacZ) (Fig. S5A) and performed Repli-seq (Fig. S4A). Of note, the converse experiment, that is, depletion of RIF1 in shMcm6 cells was precluded by the fact that the viability of shMcm6 cells was severely compromised upon viral infection with *Rif1*-specific sgRNAs or shRNAs, in line with the observation that cells with limiting MCM proteins are sensitive to stress^50–52^.

Repli-seq revealed that there was a further loss of early and late RT domain distinction in *Rif1^-/-^* shMcm6 cells compared to *Rif1^-/-^* shLacZ cells or shMcm6 cells, with a shift of both E and L RT values towards zero (Fig. 5A). An HMM-based classification of RT values, as in Fig. 1D, showed that E bins replicated later, and L bins replicated earlier in *Rif1^-/-^* shMcm6 compared to shMcm6 cells (Fig. 5B). Direct comparison of RT values in 20 kb genomic bins showed that this exacerbation of the RT phenotype in *Rif1^-/-^* shMcm6 relative to *Rif1^-/-^* shLacZ and shMcm6 cells was observed in most of the RT bins (Fig. 5C). Additionally, in *Rif1*^-/-^ shMcm6 cells, many large early-replicating domains showed considerable fluctuation in the RT values resulting in a highly fragmented RT profile (Fig. 5D).

**Figure 5:**
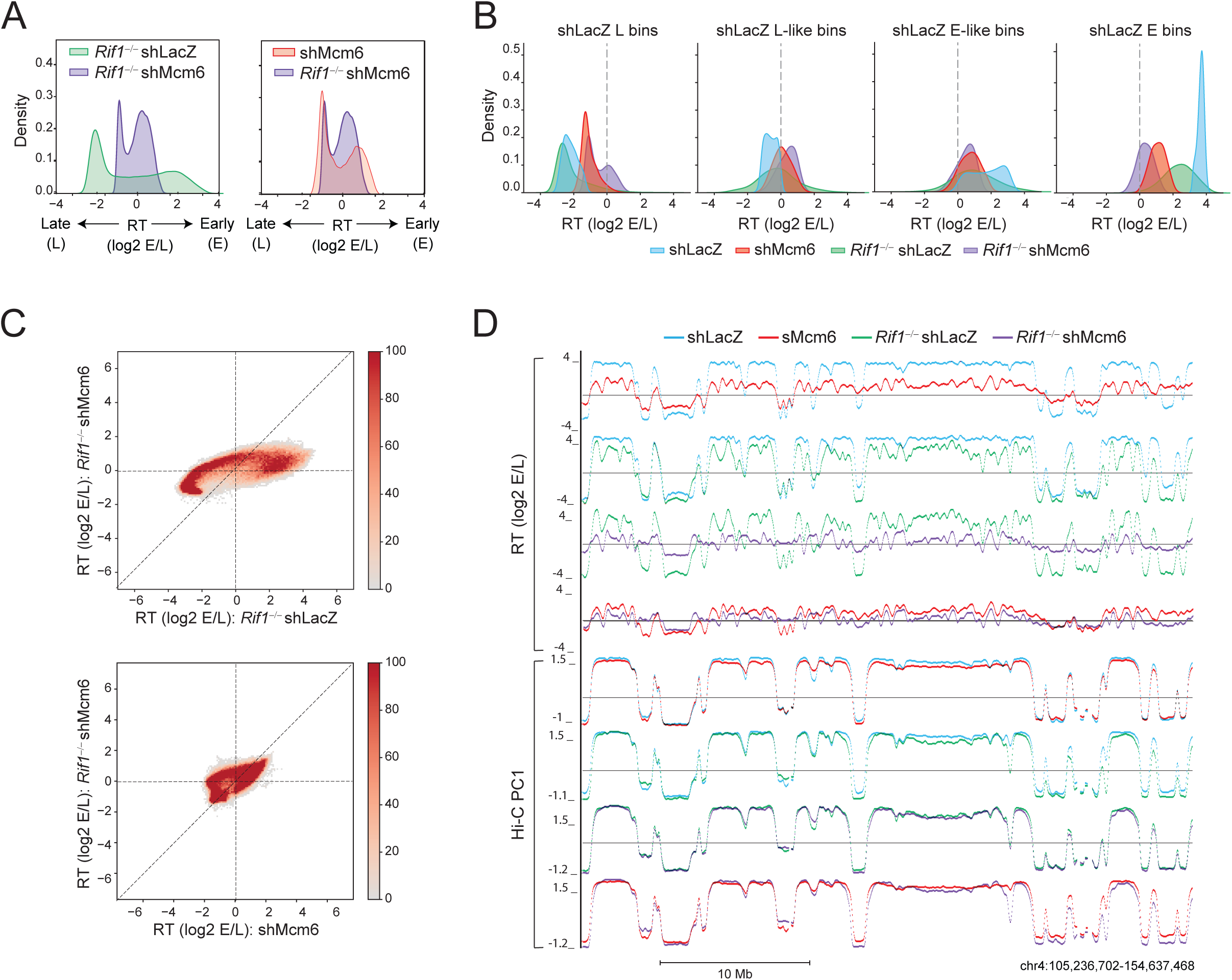
RIF1 and MCM complexes act in an additive manner to regulate early RT in B cells. A. RT histogram comparing *Rif1*^-/-^ shLacZ and *Rif1*^-/-^ shMcm6 CH12 cells. B. Identical to Fig. 1G showing how HMM-based RT states called in shLacZ cells change in the other three conditions. C. Comparison of RT values in 20 kb genomic bins between *Rif1*^-/-^ shLacZ and *Rif1*^-/-^ shMcm6 cells (top) and between shMcm6 and *Rif1*^-/-^ shMcm6 cells (bottom). D. A representative UCSC genome browser view comparing Hi-C PC1 profiles with RT (log2 E/L) profiles in shLacZ, shMcm6, *Rif1*^-/-^ shLacZ and *Rif1*^-/-^ shMcm6 CH12 cells.

However, this fragmentation was considerably lower in large late-replicating domains (Fig. 5D). The RT profiles in *Rif1*^-/-^ shMcm6 cells were also marked by switching of RT signatures (both E to L and L to E) and loss of clear domain boundaries (Fig. 5D). These changes, and especially the global fluctuations of early RT values, result in *Rif1*^-/-^ shMcm6 cells acquiring an RT signature distinct from shMcm6 cells (Fig. 5A, C). In sum, these findings reveal an additive effect of MCM depletion and loss of RIF1 on RT in B cells.

Despite the changes in RT, gene expression (Fig. S4B-C and Table S2), compartment identity (Fig. 5D and Fig. S4D) and compartment strength (Fig. S4E-F) were not majorly altered in *Rif1*^-/-^ shMcm6 cells relative to *Rif1^-/-^* shLacZ cells, although slight gains in A-B, B-A and A-A interactions were observed, akin to the changes seen in shMcm6 cells (Fig. S4F, compare with Fig. 2F). However, the major changes in RT led to a distinct pattern of PC1 versus RT profiles in in *Rif1*^-/-^ shMcm6 cells relative to in *Rif1*^-/-^ shLacZ cells (Fig. S4G). Thus, the deregulation of the RT program by MCM depletion does not majorly impact genome architecture in normal (Fig. 2) or in *Rif1*^-/-^ cells.

**Table 2:**
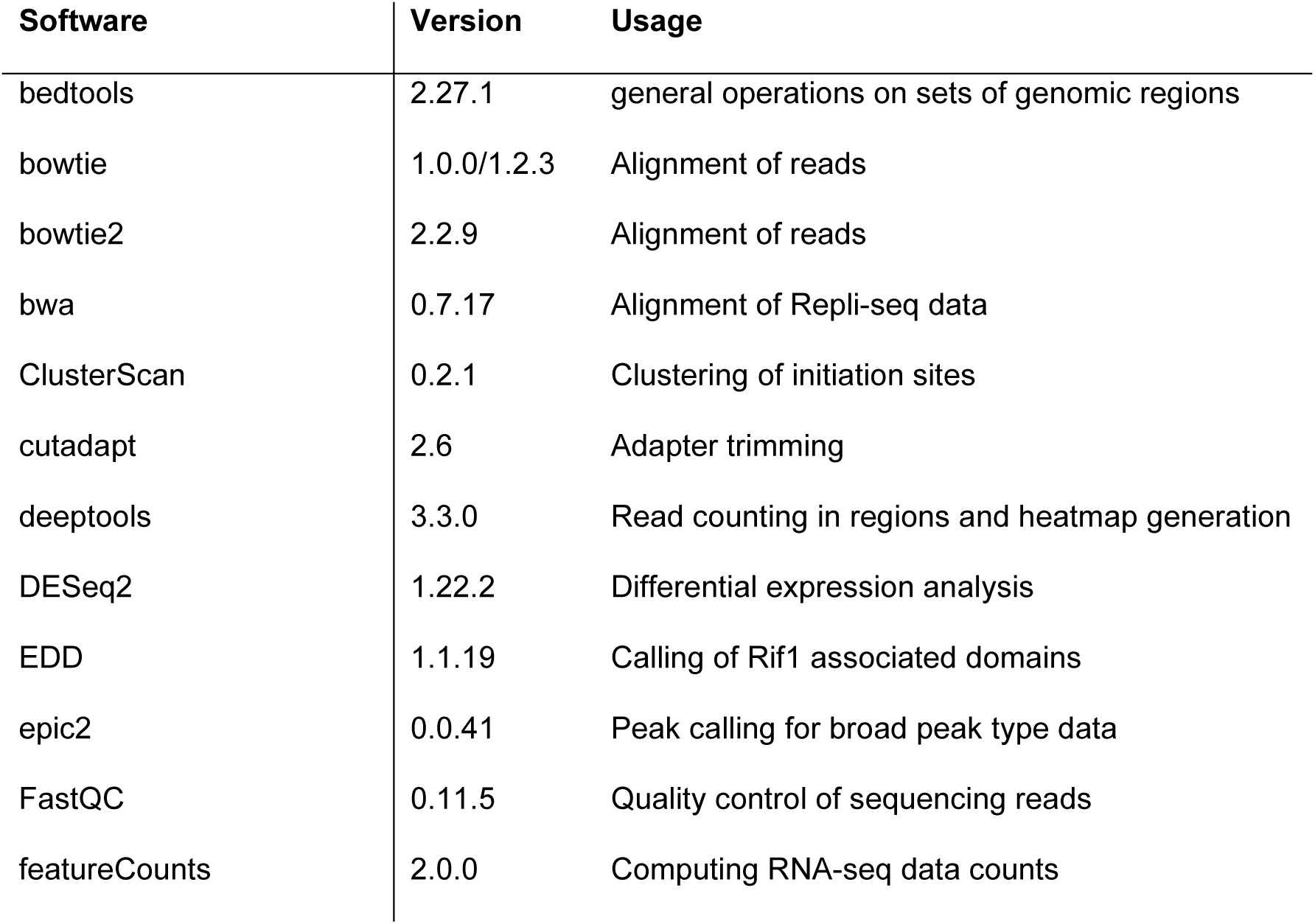
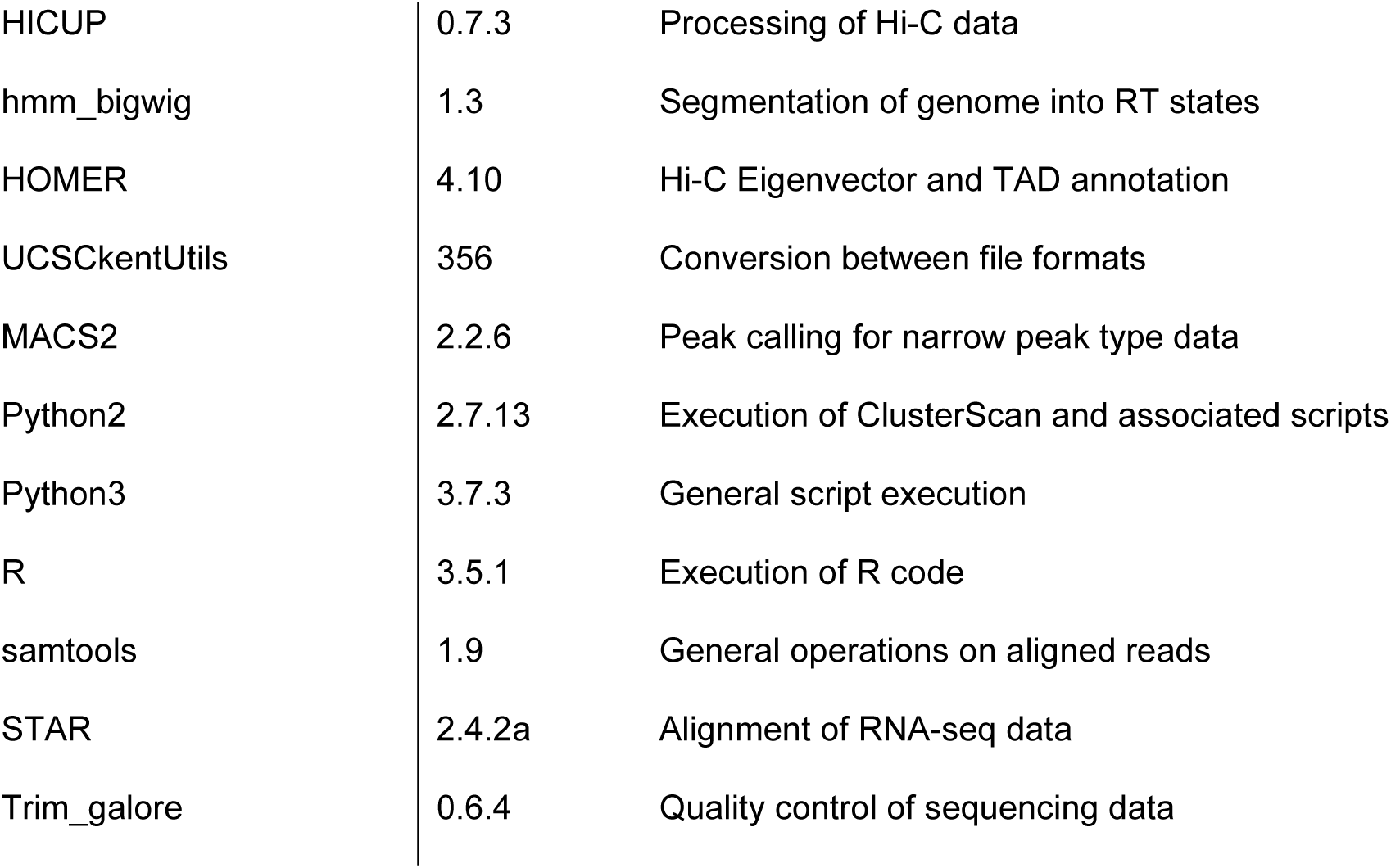
Summary of standalone software.

| Software | Version | Usage |
| --- | --- | --- |
| bedtools | 2.27.1 | general operations on sets of genomic regions |
| bowtie | 1.0.0/1.2.3 | Alignment of reads |
| bowtie2 | 2.2.9 | Alignment of reads |
| bwa | 0.7.17 | Alignment of Repli-seq data |
| ClusterScan | 0.2.1 | Clustering of initiation sites |
| cutadapt | 2.6 | Adapter trimming |
| deeptools | 3.3.0 | Read counting in regions and heatmap generation |
| DESeq2 | 1.22.2 | Differential expression analysis |
| EDD | 1.1.19 | Calling of Rif1 associated domains |
| epic2 | 0.0.41 | Peak calling for broad peak type data |
| FastQC | 0.11.5 | Quality control of sequencing reads |
| featureCounts | 2.0.0 | Computing RNA-seq data counts |
| HICUP | 0.7.3 | Processing of Hi-C data |
| hmm_bigwig | 1.3 | Segmentation of genome into RT states |
| HOMER | 4.10 | Hi-C Eigenvector and TAD annotation |
| UCSCkentUtils | 356 | Conversion between file formats |
| MACS2 | 2.2.6 | Peak calling for narrow peak type data |
| Python2 | 2.7.13 | Execution of ClusterScan and associated scripts |
| Python3 | 3.7.3 | General script execution |
| R | 3.5.1 | Execution of R code |
| samtools | 1.9 | General operations on aligned reads |
| STAR | 2.4.2a | Alignment of RNA-seq data |
| Trim_galore | 0.6.4 | Quality control of sequencing data |

To determine whether the increased deregulation of RT in *Rif1*^-/-^ shMcm6 cells was associated with changes in underlying replication origin activity, we performed SNS-seq in *Rif1*^-/-^ shLacZ and *Rif1*^-/-^ shMcm6 cells (Fig. 6A). We further classified ISs based on fold-changes (*Rif1^-/-^* shLacZ/*Rif1^-/-^* shMcm6) into quintiles and analyzed the distribution of IS densities (Fig. 6B) and RT values (Fig. 6C) within the IS classes. The most downregulated ISs (class 1) were the most active in *Rif1^-/-^* shLacZ and were predominantly early-replicating, whereas the most upregulated ISs (class 5) were the least active in *Rif1^-/-^* shMcm6 cells and were mostly late-replicating (Fig. 6B-C). This is consistent with the phenotype we reported previously in shMcm6 cells^13^. Indeed, SNS-seq profiles showed a characteristic downregulation of origin activity in A compartments accompanied by upregulation in B compartments in both shMcm6 and *Rif1^-/-^* shMcm6 cells (Fig. 6D). We infer that the additional delay in early replication in *Rif1^-/-^* shMcm6 cells relative to *Rif1^-/-^* shLacZ cells is likely due to the further decrease in the efficiency of early-firing origins.

**Figure 6:**
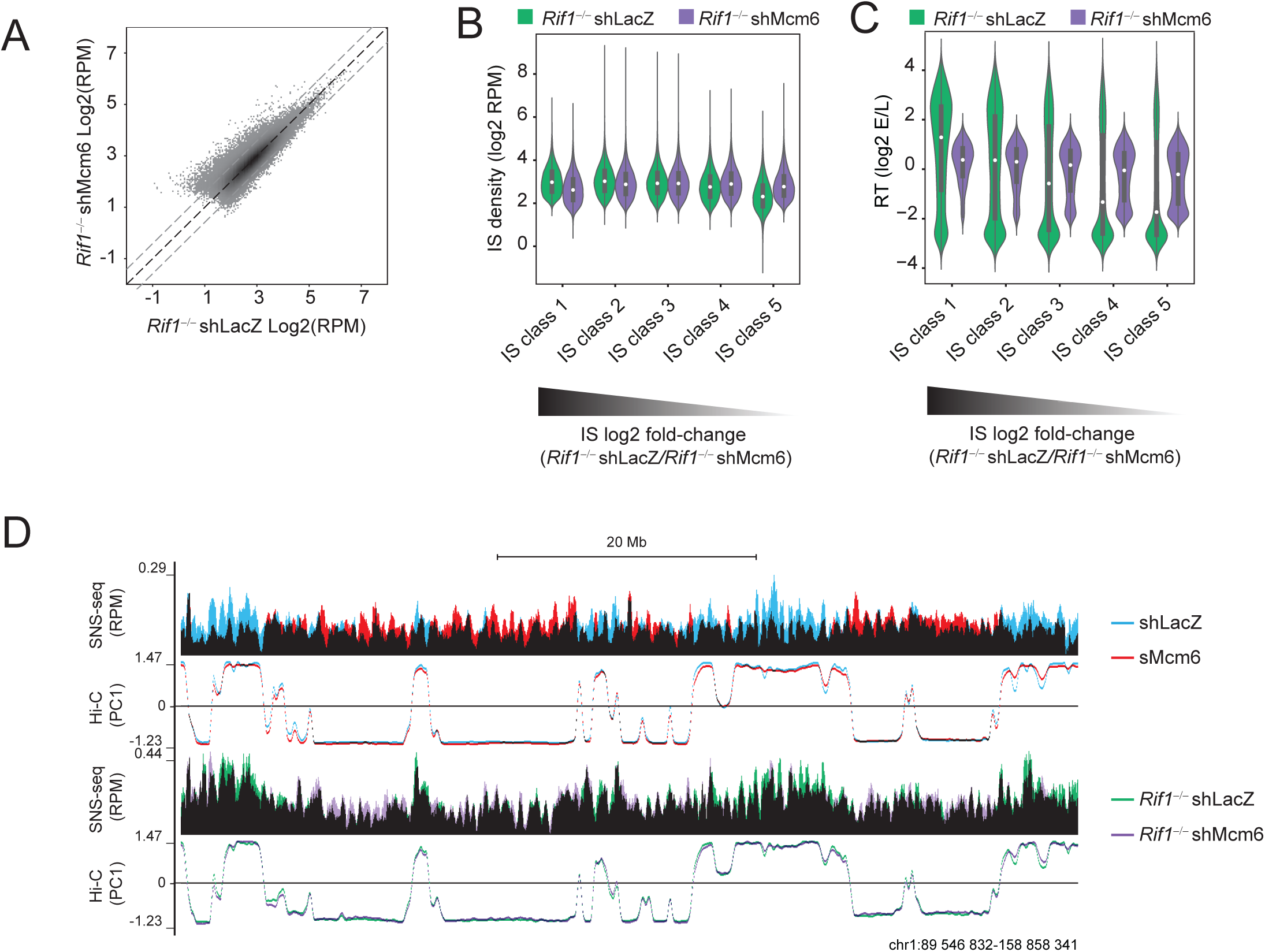
MCM depletion in *Rif1***^-/-^** cells leads to a further deregulation of early origin activity. A. Scatter plot of IS density in log2 RPM in *Rif1*^-/-^ shLacZ and *Rif1*^-/-^ shMcm6 cells identified from SNS-seq. B. ISs were split into five equal classes (quintiles) based on their fold-change (log2 *Rif1*^-/-^ shMcm6/ *Rif1*^-/-^ shLacZ) such that class 1 contained the most downregulated ISs and class 5 contained the most upregulated ISs, respectively, in *Rif1*^-/-^ shMcm6 cells. The violin plots show the IS density within each class in *Rif1*^-/-^ shLacZ and *Rif1*^-/-^ shMcm6 cells. C. ISs were classified into five classes as in C above. The violin plots show the RT values within each IS class. D. Representative genomic snapshot of SNS-seq and HiC PC1 profiles. Data from shLacZ (blue) and shMcm6 (red) are overlaid with black being the overlap between them. *Rif1*^-/-^ shLacZ and *Rif1*^-/-^ shMcm6 tracks are in green and purple, respectively, with black indicating the overlap between them.

We conclude that MCM proteins and RIF1 regulate RT in a complementary and non-epistatic manner in B cells with the contribution of each factor correlating with their impact on origin firing efficiency.

### Within early RT domains, the RT of highly transcribed regions is most sensitive to the depletion of MCM complexes and, to a weaker extent, the loss of RIF1

A closer examination of the RT profiles revealed that that the extensive fragmentation seen in *Rif1*^-/-^ shMcm6 cells was also visible to a lesser extent in shMcm6 and to the weakest extent in *Rif1*^-/-^ shLacZ cells (Fig. 5C-D). Furthermore, we observed a high degree of similarity in the fragmentation patterns between the different conditions. This suggested that these were not random fluctuations of the RT signals but reflected an underlying mechanism supporting early replication that is reliant on MCM proteins and RIF1. We therefore investigated whether the levels of nascent transcription within genomic RT bins could explain these altered RT profiles. This reasoning was based on the observations that fragmentation was typically seen in early RT domains where most transcribed genes are located, and secondly, that in *Rif1*^-/-^, shMcm6 and *Rif1*^-/-^ shMcm6 cells, early origins in active chromatin were downregulated despite there being no major changes in transcription.

To address this, we divided the genome into 20 kb bins and extracted all early-replicating (E) bins in shLacZ cells using the HMM-based approach described in Fig. 1C. Within this group, we defined transcribed bins as those having a PRO-seq density (RPM) ≥ 10 and these were divided into three equal sub-groups (tertiles) based on their RPM values, termed High (199-3358 RPM), Medium (78-198.9 RPM) and Low (10-77.9 RPM) (Fig. S4H). The remaining early-replicating bins were termed Untranscribed (0-9.9 RPM) (Fig. S4H). To compare changes in RT values, we generated RT heatmaps for the four transcription-based groups and ranked each of them by decreasing WT PRO-seq read density such that, effectively, the entire set of bins were ranked from highest to lowest PRO-seq density in shLacZ cells (Fig. 7A). In addition, we generated density plots to visualize the differences in the distribution of RT values between the four groups in all experimental conditions (Fig. 7B).

**Figure 7:**
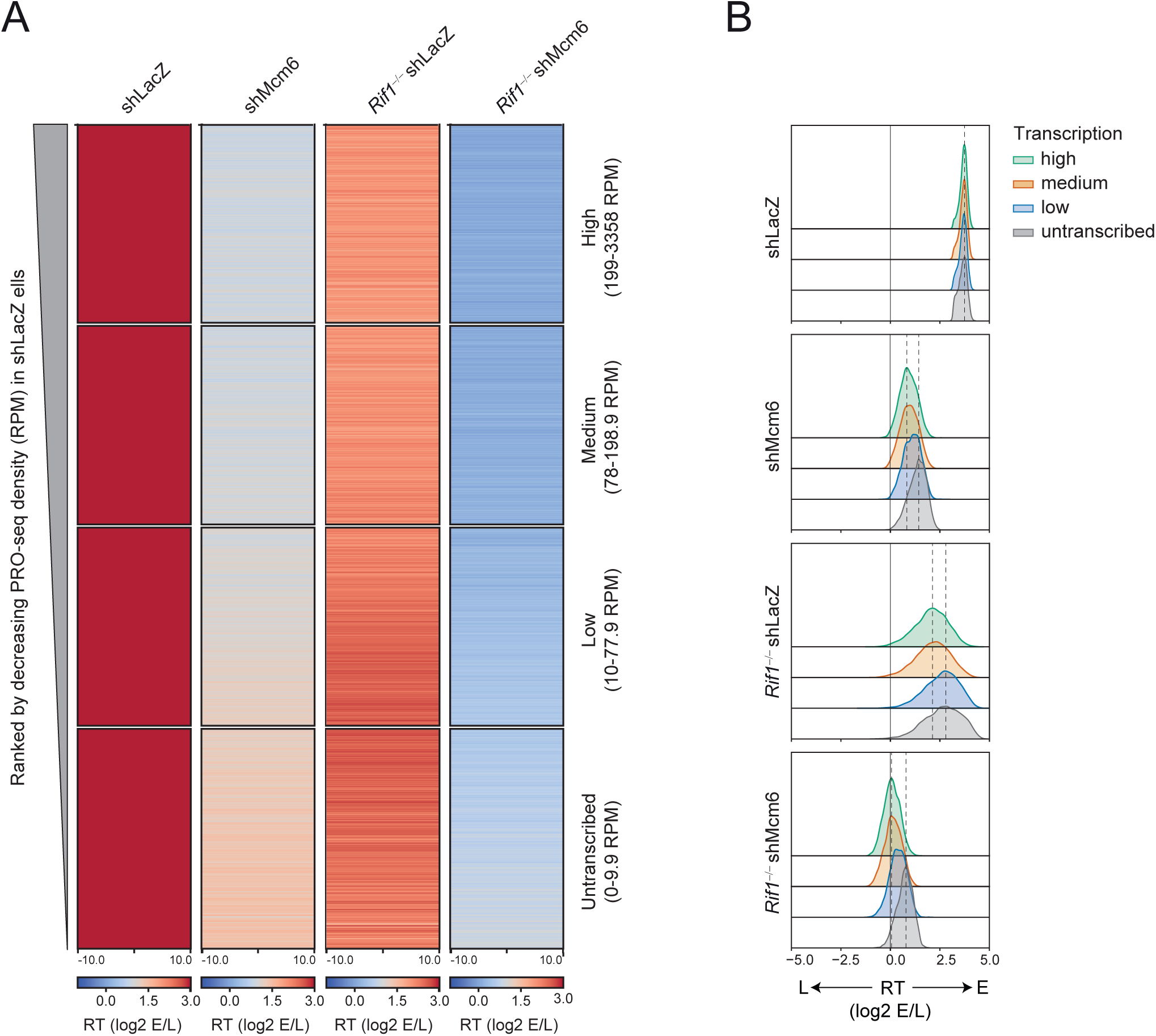
Within early-replicating domains, the RT of highly transcribed regions is more sensitive to reduction in MCM proteins and loss of RIF1 than poorly transcribed regions. A. Heatmap of RT values in 20 kb E bins from shLacZ cells (from Fig. 1G) that were split into four groups based on PRO-seq RPM. The range of PRO-seq RPMs in each group is indicated on the left. All heatmaps are ranked from highest to lowest PRO-seq density such that the entire set of heatmaps is effectively ranked from highest to lowest PRO-seq density for all shLacZ E bins. B. Density distribution of the RT values in the four transcription-based groups from A above. The dashed black lines indicate the modes of the distributions in the High (green) and Untranscribed (grey) classes.

In shLacZ cells, all four transcription-based groups showed nearly identical distributions of RT values, indicating that transcribed and non-transcribed regions normally have similar probabilities of origin activation and early replication (Fig. A-B). The High and Medium groups appeared very similar in terms of the RT values between bins (Fig. 7A) and the overall distribution profiles (Fig. 7B). However, in shMcm6 and *Rif1*^-/-^ shMcm6 cells, the High and Medium transcribed bins showed the strongest delays in RT whereas the Untranscribed group showed the least delays (Fig. 7A-B). These differences were observed across all bins (Fig. 7A). The shift in RT between the High and Untranscribed groups was ∼2-fold (log2 RT ∼1), as measured by the distance between the modes of the distributions (marked by dotted lines in Fig. 7B). The lowly transcribed regions had an intermediate distribution of RT values between the High/Medium and Untranscribed groups in both shMcm6 and *Rif1*^-/-^ shMcm6 cells (Fig. 7A-B). Thus, the strongest delays in RT are associated with regions harboring higher levels of transcription. In *Rif1*^-/-^ shLacZ cells, all groups had a broad spread of RT values with similar distribution profiles (Fig. 7A-B). However, a closer inspection of the heatmap and the modes of the distributions revealed a slightly greater delay in RT for the high and medium transcribed regions compared to the lowly transcribed and non-transcribed bins (Fig. 7B).

Taken together, these results suggest that the RT of transcribed regions, and hence the underlying activity of early origins, is more sensitive to the reduction of MCM proteins than to the loss of RIF1, which correlates with the magnitude of the fragmentation observed in the early RT domains (Fig. 5C-D). We conclude that the fragmented appearance of early RT profiles is due to underlying transcription, with stronger delays correlating with higher levels of transcription. These results suggest that transcriptional strength negatively regulates the probability of early origin firing at active genes, and that this is normally overcome by the presence of a normal complement of MCM proteins and, to a lesser extent, RIF1.

## DISCUSSION

Our study reveals two new layers of regulation within the RT program. First, a mechanism that specifically promotes early RT via the functional repurposing of RIF1. Second, the importance of normal levels of MCM proteins, with a minor role for RIF1, in ensuring early RT of transcribed regions. We propose that since mutation rates are known to be higher in late-replicating regions ^11, 14–17^ and delays in RT are the major cause of common fragile sites in long transcribed genes^18–20^, ensuring early RT may be a safeguarding mechanism to ensure genome integrity and normal cellular functions post mitosis. As explained below, this may be particularly relevant for antigen-activated B cells.

How might transcription negatively impact the efficiency of replication origins? There is evidence from *in vitro* studies that RNA polymerase (RNAP) complexes can push MCM complexes along DNA ^53, 54^. Indeed, MCM and ORC proteins were found to be relatively depleted in gene bodies^55^ and Okazaki fragment sequencing (OK-seq) studies have reported that replication initiation zones occur largely in the intergenic regions flanking transcribed genes^56^. A recent single molecule imaging study, where replication-transcription encounters were reconstituted using purified proteins, may be instructive in this regard ^54^. This study found that T7 RNAP could efficiently push DNA-bound ORC, OCCM (an intermediate of the pre-RC consisting of ORC, Cdc6, Cdt1 and a single MCM hexamer) and MCM double hexamers (the configuration within the fully assembled pre-RC). However, whereas OCCM and double hexamers were rarely ejected by RNAP, ORC alone was frequently evicted by RNAP. Moreover, most of the ORC molecules repositioned by RNAP were unstable ^54^. Since ORC loading is the first step in origin licensing ^25^, the inference is that the labile binding of ORC makes licensing in transcribed regions inherently less efficient than in non-transcribed regions, but that this is overcome by the association of MCM complexes, which minimizes the loss of single ORC complexes by RNAP.

Importantly, our data show that, normally, the distribution of early RT values is similar between transcribed and non-transcribed regions. Hence, we propose that, in WT cells, the large pool of MCM complexes ensures that the loading of ORC is rapidly followed by the assembly of the OCCM and pre-RC such that the loss of single ORC complexes by RNAP is minimized. This ensures that both transcribed and non-transcribed regions have a similar probability of early origin activation and early replication in WT cells. When MCM complexes are limiting, licensing in both transcribed and non-transcribed regions is reduced. However, given the inhibitory effect of transcription on ORC stability, the formation of OCCM and pre-RC formation in highly transcribed regions will be inefficient, allowing for more eviction of ORC by RNAP. This will reduce early origin licensing efficiencies in highly transcribed chromatin to a greater degree than in non-transcribed chromatin, leading to our observation of a higher probability of early replication in non-transcribed regions relative to transcribed regions.

Although the mechanism by which RIF1 regulates early origin firing in B cells remains to be deciphered, RIF1 can interact with MCM complexes in activated B cells ^38^. Intriguingly, a previous study in human cells reported a role for RIF1 in promoting origin licensing by protecting ORC1 from phosphorylation-mediated degradation ^57^ . Thus, it is plausible that RIF1 prevents premature disassembly of OCCM and pre-RC complexes in B cells. In addition, although RIF1 binds broadly across early-replicating chromatin domains in B cells, ∼70% of RIF1 peaks and ∼50% of RADs in B cells are in genes (Fig. 3). This suggests that the protective function of RIF1 in origin licensing may be more pronounced in highly transcribed regions. According to this model, the loss of RIF1 would lead to fewer licensed origins in highly transcribed regions compared to neighboring non-transcribed regions within the same early-replicating domains, resulting in non-transcribed regions replicating slightly later than highly transcribed regions, as we observe.

Given that RIF1 has not been previously implicated in regulating early replication, our study raises the question of why antigen-activated B cells have functionally repurposed RIF1 in this manner. Antigen-activated B cells are amongst the fastest proliferating cells in the body with a cell cycle duration of 6-8 h *in vivo* within germinal centers and 8-12 h in culture^58–60^. A unique feature of these cells is that they undergo extremely high rates of somatic hypermutation at the highly transcribed immunoglobulin genes, a necessary event in antibody maturation upon infection or vaccination.

Genome instability is further elevated by the fact that somatic hypermutation also occurs at many other transcribed loci, including proto-oncogenes like *BCL6* and *MYC* ^61–64^, which can result in oncogenic translocations typical of most mature B cell cancers ^65–67^. Moreover, the DNA repair pathways activated by somatic hypermutation result in single- and double-strand breaks as well as singe-strand patches, all of which are impediments for replication^68, 69^. Therefore, these B cells proliferate in a highly genotoxic environment where replication stress is elevated. Under such conditions, it is plausible that B cells have functionally repurposed RIF1 as an additional mechanism to safeguard its genome by enforcing the early replication of active genes, which would help to decrease mutations and common fragile sites associated with late replication or delayed replication of long genes^11, 14–20^. It is also possible that this function of RIF1 is masked in other cell types where the dominant mode of RT regulation is that of RIF1-mediated suppression of origins in late-replicating chromatin^7, 34, 35^.

## METHODS

### Cell culture

CH12 cells were maintained in RPMI medium with 10% fetal bovine serum (FBS), glutamine, sodium pyruvate, Hepes and antibiotic/antimycotic mix. LentiX packaging cells were maintained in DMEM medium with 10% FBS and antibiotics.

### Transfections and infections

Transfection of LentiX cells with shRNAs against LacZ and Mcm6 followed by infection of CH12 cells with lentiviral supernatants was done exactly as described in our previous study^39^.

### Sample preparation for replication timing (RT) analysis

Repli-seq was performed as previously described^40^. In brief, two million asynchronously dividing cells were seeded and incubated with 100μM BrdU (Sigma) for 2h in a light protected environment to maintain BrdU stability. Cells were fixed and incubated with a mix of RNase A (Invitrogen) and propidium iodide (Sigma) for 30 min (light protected). For each sample, three fractions were sorted: G1 phase, early S phase and late S phase cells, and for each fraction, two independent samples of 50,000 cells (technical replicates) were sorted on a Sony SH800S Cell Sorter. Sorted cells were lysed with Proteinase K buffer overnight. Extracted DNA was sonicated for 9 min in a Diagenode Bioruptor resulting in 100-500 bp DNA fragments as determined on an agarose gel. Sonicated DNA was subjected to end-repair and adapter ligation using the NEBNext® Ultra™ II DNA Library Prep Kit (NEB) following the NEB protocol. Adapter-ligated DNA was incubated with 25 μg/ml of anti-BrdU antibody (BD Pharmingen) for 4h with rotation followed by incubation with 40 μg of anti-mouse IgG antibody (Sigma) for 1h with rotation (light protected). DNA was precipitated via Centrifugation at 16,000*g* for 5 min at 4°C. Pellet was resuspended in 200 μl of digestion buffer( for 50 ml of digestion buffer, combine 44 ml of autoclaved double-distilled water, 2.5 ml of 1M Tris-HCl, pH 8.0, 1 ml of 0.5M EDTA and 2.5 ml of 10% SDS) with freshly added 0.25 mg/ml proteinase K and incubated overnight at 37°C. The immunoprecipitated DNA was used for Repli-qPCR or next generation sequencing (Repli-seq). Libraries that were successfully validated by Repli-qPCR were sequenced on an Illumina HiSeq 2500 machine (50bp, single-end). Up to 12 barcoded samples were pooled per lane.

### Chromatin immunoprecipitation for RIF1 occupancy in primary B cells

Chromatin immunoprecipitation was performed as described ^36^. In summary, primary B cells were isolated from Rif1^FH/FH^ and C57BL/6 (Rif1^WT/WT^) mice spleens using anti-CD43 MicroBeads (Miltenyi Biotec) and expanded in complete RPMI containing 5 µg/ml LPS (Sigma-Aldrich) and 5 ng/ml mouse recombinant IL-4 (Sigma-Aldrich) to allow B cell activation and class switch recombination to IgG1. B cells were harvested 72h after activation and 4 x 10^7^ cells were cross-linked by using 2 mM disuccinimidyl glutarate (ThermoFisher 20593) in PBS for 45 min and 1% formaldehyde for 5 min (Thermo Scientific 28908). The reaction was quenched with 0.125 M glycine. Cells were washed thrice with ice-cold PBS and lysed in SDS lysis buffer. Chromatin fragmentation was performed using a Covaris E220 sonicator to obtain fragments between 200 and 600 bp. Chromatin was quantified with a ND-1000 NanoDrop spectrophotometer. Immunoprecipitation was performed with 0.5 μg of anti-HA antibody (Santa Cruz sc-7392) and 50 µl of Dynabeads protein G (Thermofisher 10003D) and 25 μg chromatin. ChIP libraries were prepared by NEBNext ultra II DNA library preparation kit (NEB E7645L) and sequenced on one lane of a NovaSeq 6000 (Illumina) machine.

### RNA-seq from CH12 cells

RNA-seq was performed on CH12 and two independent RIF1-deficient (RIF1^-/-^) CH12 clonal derivatives ^37^ with three replicates. Cells were expanded in complete RPMI and 1 million cells were collected by centrifugation. RNA was extracted with TRIzol (Invitrogen) according to manufacturer’s instructions. TruSeq RNA Library Prep Kit v2 (Illumina) was used to prepare a whole-transcriptome sequencing library and sequenced on one lane of a NovaSeq 6000 SP (Illumina) machine.

### Isolation of short nascent strands (SNSs)

SNSs were isolated following an established protocol^48^ kindly provided by Dr. Maria Gomez (CBMSO, Madrid) and also described in our previous study (Wiedemann et al, 2016). In brief, 200 million cells per replicate from asynchronous cell cultures were harvested and genomic DNA was extracted. DNA was denatured at 95°C for 10 minutes and then subjected to size fractionation via 5-20% neutral sucrose gradient centrifugation (24,000g for 20h). Fractions were analyzed by alkaline agarose gel electrophoresis and those in the 500-2000 nt range were pooled. Prior to all following enzymatic treatments, ssDNA was heat denatured for 5 min at 95°C. DNA was phosphorylated for 1h at 37°C with T4 Polynucleotide kinase (NEB or made in-house by the Molecular Biology Service, IMP). To enrich for nascent DNA strands, the phosphorylated ssDNA was digested overnight at 37°C with Lambda exonuclease (NEB or made in-house by the Molecular Biology Service, IMP). Both T4 PNK and Lambda exonuclease steps were repeated twice for a total of three rounds of phosphorylation and digestion. After the final round of digestion, the DNA was treated with RNaseA/T1 mix (Thermo Scientific) to remove 5’ RNA primers and genomic RNA contamination. DNA was purified via phenol-chloroform extraction and ethanol precipitation This material was either used directly for qPCR or further processed for library preparation.

### SNS library preparation for SNS-seq

SNSs prepared as described above were converted to double stranded DNA (dsDNA) via random priming with random hexamer primer phosphate (Roche) and ligation with Taq DNA ligase (NEB)^48^. DNA was checked on a fragment analyzer and 50ng was used for library preparation with the NEBNext® Ultra™ II DNA Library Prep Kit (NEB) following the manufacturer’s protocol. Libraries were barcoded using the NEBNext® Singleplex Oligos (NEB) as per the NEB protocol which allowed 4-8 libraries to be pooled per run. Sequencing was performed on an Illumina HiSeq 2500 machine (50 bp, single-end).

### In situ Hi-C

Hi-C was performed as previously described^43^ with minor modifications. In brief, 5 million cells were crosslinked with 1% formaldehyde (Sigma) for 10 min and quenched with 0.6M glycine for 5 min. Cells were lysed with Hi-C lysis buffer for 1h on ice and nuclei were collected by centrifugation. Nuclei were digested with 375U of Mbol (NEB) overnight at 37°C with rotation. Biotin-14-dATP (Life Technologies) was incorporated for 1h at 37°C with rotation. Ligation of overhangs was performed with 20,000U of T4 DNA ligase (NEB) for 4h at room temperature with rotation. Nuclei were pelleted and reverse-crosslinked overnight. Purified DNA was sonicated for 14 min in a Diagenode Bioruptor to obtain a size range of 200-700bp. This material was purified using Agencourt AMPure XP beads (Beckman Coulter). Between 8-15 μg of DNA was incubated with 100μl (∼10 mg) of Dynabeads MyOne Streptavidin C1 (Invitrogen) for 15 min with rotation. End repair and adapter ligation using the NEBNext® Ultra™ II DNA Library Prep Kit was performed on-beads following the kit manual. The adapter-ligated DNA was washed, eluted and PCR-amplified with KAPA 2X HiFi HotStart ReadyMix (Kapa Biosystems) and the NEBNext Multiplex Oligos for Illumina® (Dual Index Primers Set 1). Four pooled, barcoded samples were sequenced on an Illumina NovaSeq 6000 machine (50 bp, paired-end).

### PRO-seq

PRO-seq was performed as described previously^70^ with minor modifications. To isolate nuclei, CH12 cells and *Drosphila* S2 cells were resuspended in cold Buffer IA (160mM sucrose, 10mM Tris-Cl pH 8, 3mM CaCl2, 2mM MgAc2, 0.5% NP-40, 1mM DTT added fresh), incubated on ice for 3 min and centrifuged at 700g for 5 min. The pellet was resuspended in nuclei resuspension buffer NRB (50mM Tris-Cl pH 8, 40% glycerol, 5mM MgCl2, 0.1mM EDTA). For each run-on, 10 million CH12 nuclei were spiked with 10% *Drosophila* S2 nuclei in a total of 100µL NRB and incubated at 30°C for 3 min with 100 µL 2x NRO buffer including 5µl of each 1mM NTP (biotinylated ATP and GTP, and unlabelled UTP and CTP). The following steps were performed as described^70^ with the following changes: (1) we used different adapters, namely, 3 RNA adapte 5Phos/NNNNNNNGAUCGUCGGACUGUAGAACUCUGAAC/3InvdT-3’ and 5’RNA adapter: 5’-CCUUGGCACCCGAGAAUUCCANNNN-3’. (2) 3’ and 5’ ligations which were done at 16°C overnight, and (3) CapClip pyrophosphatase (Cellscript) used for 5’ decapping. RNA was reverse transcribed by SuperScript III RT (Invitrogen) with RP1 Illumina primer to generate cDNA libraries. Libraries were amplified using barcoding Illumina RPI-x primers and the universal RP1 and KAPA HiFi Real-Time PCR Library Amplification Kit. Amplified libraries were subjected to electrophoresis on 2.5% low melting agarose gel and amplicons from 150-350 bp were extracted from the gel, multiplexed and sequenced on Illumina platform NextSeq 550 SR75. Bioinformatics analyses were performed as described^70^ but additionally the random 8-mer was used to exclude PCR duplicates and only deduplicated reads were aligned.

### Biological and technical replicates

For biological replicates, different frozen vials of CH12 cells were thawed and kept separate throughout the course of the experiment. Replicates for all next-generation sequencing experiments were derived in this manner. Technical replicates, where used (such as in RT-qPCR or ChIP-qPCR), were subsets of the biological replicate.

### Statistical analyses

For correlation scatter plots, the Pearson correlation coefficient was calculated to determine the degree of correlation. In all other cases, the two-tailed Student’s t test was used for statistical significance.

## Bioinformatics

Main next generation sequencing data analysis workflows for SNS-seq, Repli-seq and Hi-C have been wrapped with *Nextflow* workflow language ^71^ and are centrally available at https://github.com/pavrilab ^13^. The usage of Docker and Singlularity containers ensures portability, reproducibility and reliability for all workflows. Along with integration of GitHub repositories for self- contained pipelines all workflows are easily ported to all major HPC computing platforms such as SGE, SLURM, AWS. Furthermore, continuous checkpoints for pipeline execution allow for resuming and automatic retrial of failed steps. All workflows produce elaborate QC-reports and out-of-the box, resources consumption reports to allow tailoring resources requirements to your datasets which especially for Hi-C datasets can vary by several orders of magnitudes.

Processing of raw Repli-seq, Hi-C, SNS-seq and RNA-seq data was done as described previously.^13^

### Generating heatmaps for IS peak summit neighborhoods

Sequencing coverage normalized bigWig tracks were generated from raw aligned reads of NGS data sets with deeptools bamCoverage v3.3.0 ^72^ using command-line parameters --normalizeUsing CPM -- exactScaling and --ignoreDuplicates. Next, we computed the position of the replication start site (RSS) as the genomic coordinate of the maximum pile-up of the merged tracks of compared conditions. The generated summit positions were then grouped by their associated peak class and sorted in decending order on the log2-ratio of CPM values of treatment versus control. The sorted summits were then used as reference points for deeptools computeMatrix v3.3.0 ^72^ to compute signal distributions for previously generated CPM-normalized bigWig tracks within a 4kb region centered on the summit using a binsize of 50 bp and setting missing values to zero. The results were then plotted using the associated plotHeatmap command.

### PRO-seq analysis

The 3’ end sequence of the reads (NNNNTGGAATTCTCGGTGCC) were removed using cutadapt v1.4.2 ^73^ and 9 nucleotides from their 5′ ends, containing the random 8mer and *in vitro* run-on nucleotide, were removed. The trimmed reads were reverse complemented using fastx_reverse_complement (http://hannonlab.cshl.edu/fastx_toolkit; version 0.0.13) followed by deduplication based on the 8mer sequence. Trimmed reads longer than 18 bp were aligned to a hybrid mouse drosophila genome (mouse genome build NCBI m37, GCA_000001635.18, drosophila FlyBase release 5) using bowtie v1.0.0 ^74^ with -v 2 --best --strata --tryhard -m 1 --chunkmbs 256. Unique mappers from the resulting BAM file were used to create bigwigs with deeptools bamCoverage 3.3.0.

### RT domain analysis

20kb binned RT tracks for all conditions were segmented into three states using hmm_bigwigs (https://github.com/gspracklin/hmm_bigwigs). Fitted states were then mapped to either early (E), mid (M) or late (L) replication timing based on the RT value distribution of each state (E for RT > 0, L for RT < 0 and M for RT ∼ 0). M state bins were split at 0 into L-like (values < 0) and E-like (values > 0) state bins.

### Generating saddle plots from Hi-C data

Saddle plots were computed using cooltools v0.3.2. In brief, we subdivided all bins of a 20Kb KR-normalized contact matrix into 50 equal-sized groups based on the bins compartment signal as derived from the eigenvector of the WT data, where group 1 has the lowest signal (i.e. most B) and group 50 has the highest signal (i.e. most A). Subsequently,we compute the mean observed/expected value for each pair of groups and plot it as a 50 x 50 matrix. Similarly, replication timing and H3K9me3 data can be used for bin group assignment.

### RIF1 peak and RIF1 associated domains (RADs) analyses

Peaks and domains (RADs) were called as described previously ^75^ additionally calling narrow peaks with MACS2. In brief, raw ChIP-seq and input reads in MEFs and primary B cells were trimmed with trim_galore v0.6.4 (https://github.com/FelixKrueger/TrimGalore) setting minimum length to 18 bases and default arguments otherwise and aligned to the mm9 reference genome with bowtie v1.0.0 ^74^ with -S --trim5 0 --trim3 0 -v 2 --best --strata --tryhard -m 1 --phred33-quals --chunkmbs 256. Subsequently, RIF1 narrow and broad peaks were called for each replicate with MACS v2.2.6 ^76^, RIF1 associated domains (RADs) were called using EDD v1.1.19 ^77^. RADs, narrow and broad peaks were merged using bedtools v2.27.1 ^78^ and only those peaks called in both replicates were retained for downstream analysis. Genome coverage tracks were computed with deeptools bamCoverage v3.3.0 ^72^ using command-line parameters --normalizeUsing CPM --exactScaling –binSize 50 and – ignoreDuplicates. Log2 ratio tracks were computed with deeptools bamCompare v3.3.0 ^72^ using command-line parameters --scaleFactorsMethod readCount --operation log2 --pseudocount 1 – binSize 50 –ignoreDuplicates.

### Summary of all NGS data used

**Table 1:** Summary of used data sets in this study including public Accession numbers.

| Sample Name | Data type | Source | Accession |
| --- | --- | --- | --- |
| CH12 shLacZ | SNS-seq | GEO | GSE161822 |
| CH12 shMcm6 | SNS-seq | GEO | GSE161822 |
| CH12 WT | SNS-seq | This study | - |
| CH12 Rif1<sup>-/-</sup> | SNS-seq | This study | - |
| CH12 Rif1<sup>-/-</sup> shLacZ | SNS-seq | This study | - |
| CH12 Rif1<sup>-/-</sup> shMcm6 | SNS-seq | This study | - |
| CH12 H3K4me1 | ChIP-seq | GEO/UCSC | GSM946546/wgEncodeEM002370 |
| CH12 H3K4me3 | ChIP-seq | GEO/UCSC | GSM946528/wgEncodeEM002366 |
| CH12 H3K9me3 | ChIP-seq | GEO/UCSC | GSM946548/wgEncodeEM002372 |
| CH12 H3K27ac | ChIP-seq | GEO/UCSC | GSM1000117/wgEncodeEM003167 |
| CH12 H3K36me3 | ChIP-seq | GEO/UCSC | GSM946530/wgEncodeEM002364 |
| CH12 DHS-seq | DHS-seq | GEO/UCSC | GSM1014153/wgEncodeEM003416 |
| CH12 shLacZ | Hi-C | GEO | GSE161822 |
| CH12 shMcm6 | Hi-C | GEO | GSE161822 |
| CH12 Rif1<sup>-/-</sup> shLacZ | Hi-C | This study | - |
| CH12 Rif1<sup>-/-</sup> shMcm6 | Hi-C | This study | - |
| CH12 shLacZ | Repli-seq | This study | - |
| CH12 shMcm6 | Repli-seq | This study | - |
| CH12 Rif1<sup>-/-</sup> shLacZ | Repli-seq | This study | - |
| CH12 Rif1<sup>-/-</sup> shMcm6 | Repli-seq | This study | - |
| CH12 shLacZ | RNA-seq | GEO | GSE161822 |
| CH12 shMcm6 | RNA-seq | GEO | GSE161822 |
| CH12 Rif1<sup>-/-</sup> shLacZ | RNA-seq | This study | - |
| CH12 Rif1<sup>-/-</sup> shMcm6 | RNA-seq | This study | - |
| CH12 shLacZ | PRO-seq | This study | - |
| primary B/CH12 input track | ChIP-seq | GEO | GSM2058441 |
| primary B Rif1<sup>FH/FH</sup> | ChIP-seq | This study | - |
| primary B Rif1<sup>WT/WT</sup> | ChIP-seq | This study | - |
| primary B WT Repli-seq | Repli-seq | This study | - |
| primary B Rif1<sup>-/+</sup> Repli-seq | Repli-seq | This study | - |
| primary B Rif1<sup>-/-</sup> Repli-seq | Repli-seq | This study | - |
| primary B WT ATAC-seq | ATAC-seq | GEO | GSE132029 |
| primary B WT PRO-seq | PRO-seq | This study | - |
| primary B WT H3K9me3 | ChIP-seq | This study | - |

### Summary of software used in the study

All custom Python scripts and tools, except for scripts using ClusterScan, were developed, tested and executed using Python v3.7.3. ClusterScan and related scripts were executed using Python v2.7.13. The used software and packages are summarized in Table , Table 3 and Table 4.

**Table 3:** Summary of R packages.

| **Package** | **Version** | **Usage** |
| --- | --- | --- |
| ashr | 2.2-47 | Differential expression analysis |

**Table 4:**
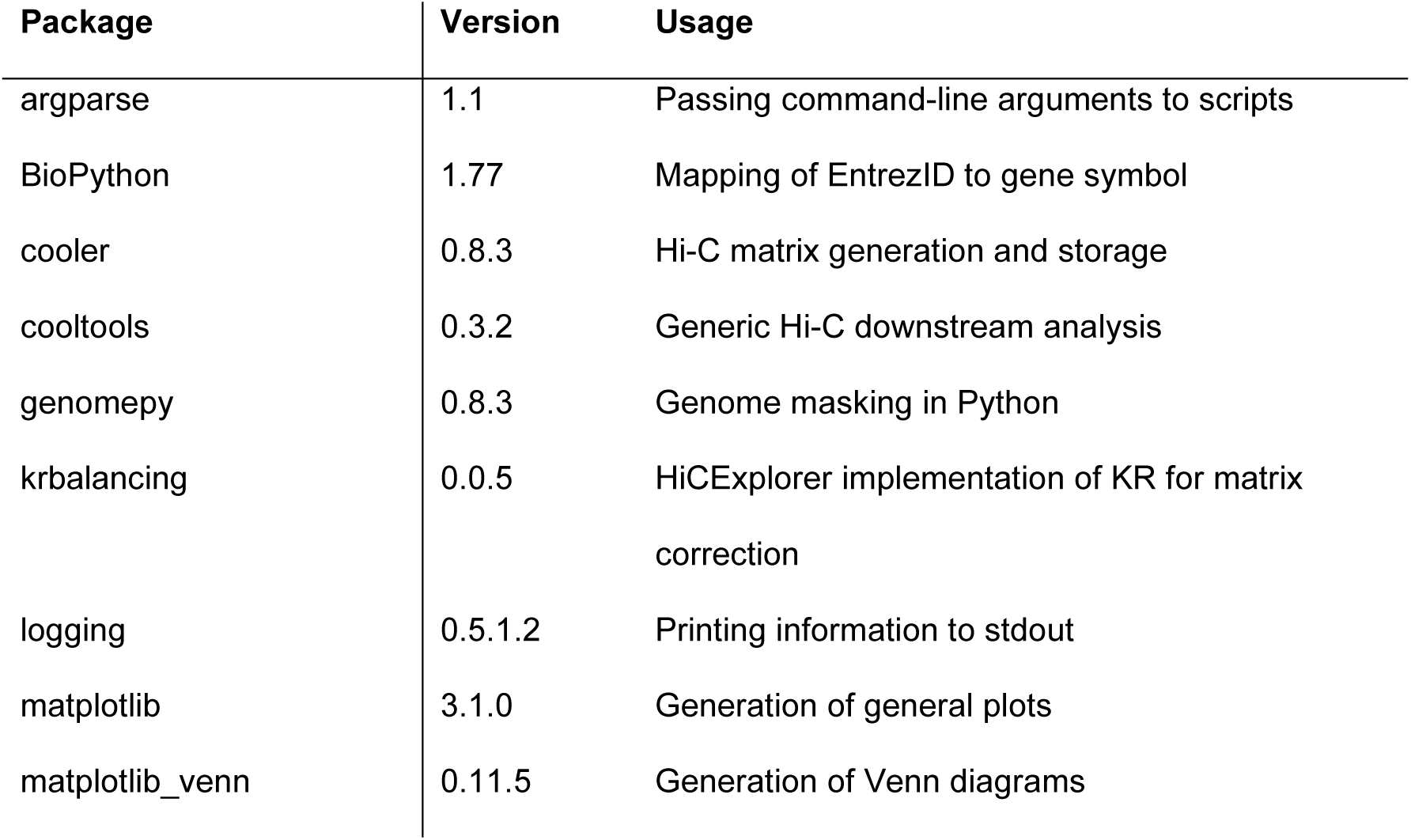
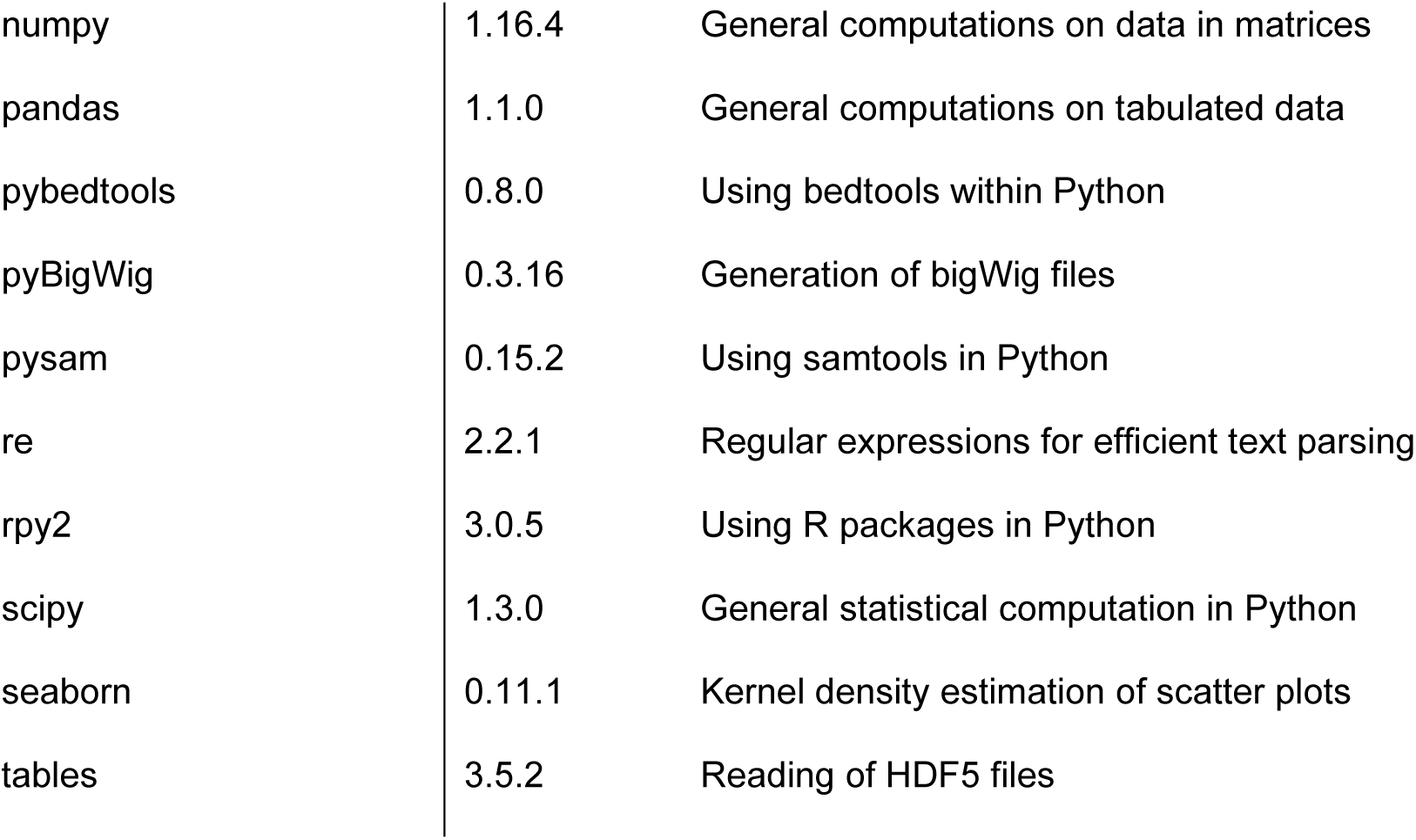
Summary of Python packages.

| **Package** | **Version** | **Usage** |
| --- | --- | --- |
| argparse | 1.1 | Passing command-line arguments to scripts |
| BioPython | 1.77 | Mapping of EntrezID to gene symbol |
| cooler | 0.8.3 | Hi-C matrix generation and storage |
| cooltools | 0.3.2 | Generic Hi-C downstream analysis |
| genomepy | 0.8.3 | Genome masking in Python |
| krbalancing | 0.0.5 | HiCExplorer implementation of KR for matrix<br>correction |
| logging | 0.5.1.2 | Printing information to stdout |
| matplotlib | 3.1.0 | Generation of general plots |
| matplotlib_venn | 0.11.5 | Generation of Venn diagrams |
| numpy | 1.16.4 | General computations on data in matrices |
| pandas | 1.1.0 | General computations on tabulated data |
| pybedtools | 0.8.0 | Using bedtools within Python |
| pyBigWig | 0.3.16 | Generation of bigWig files |
| pysam | 0.15.2 | Using samtools in Python |
| re | 2.2.1 | Regular expressions for efficient text parsing |
| rpy2 | 3.0.5 | Using R packages in Python |
| scipy | 1.3.0 | General statistical computation in Python |
| seaborn | 0.11.1 | Kernel density estimation of scatter plots |
| tables | 3.5.2 | Reading of HDF5 files |

## Data availability

All next generation sequencing data (Repli-seq, RNA-seq, Hi-C, ChIP-seq and SNS-seq) has been deposited in GEO under accession number GSE228880.

## ACKNOWLEDGEMENTS

We are grateful to the Vienna Biocenter Core Facilities (VBCF) for next-generation sequencing and the IMP/IMBA core facilities especially the animal house, bio-optics, molecular biology service and proteomics service. We are grateful to Roman Stocsits and Dr. Anton Goloborodko for advice on Hi-C data analysis. This work was funded by Boehringer Ingelheim, The Austrian Industrial Research Promotion Agency (Headquarter Grant FFG-834223), grants from the Austrian Science Fund to RP (FWF P 29163-B26) and UES (FWF T 795-B30), and the Helmholtz-Gemeinschaft Zukunftsthema "Immunology and Inflammation" ZT810 0027 to MDV and AR.

## AUTHOR CONTRIBUTIONS

DM performed most of the bioinformatics analyses, established pipelines and analyzed data. MP performed Repli-seq, Hi-C, RNA-seq and SNS-seq, and analyzed data. AR performed RIF1 ChIP-seq from primary B cells and RNA-Seq in CH12 cells. SG and KNK performed Repli-seq from primary B cells. US performed PRO-seq. MN collaborated on Repli-seq from CH12 cells. DMG, SCBB and MDV contributed personnel, resources, extensive discussions and critical reading of the manuscript. TN and RP co-supervised the project and analyzed data. RP conceived the project and wrote the manuscript with inputs from all authors.

## Supplementary Table Legends

**Table S1: RNA-seq analysis from two independent Rif1^-/-^ CH12 clones relative to WT cells**

The first two tabs contain the analysis of all genes from *Rif1*^-/-^ CH12 clones 1 and 2 analyzed using the DESeq2 software^42^. The next two tabs list the common two-fold downregulated (n = 22) and upregulated (n = 3) genes.

**Table S2: RNA-seq analysis from Rif1^-/-^ shLacZ and Rif1^-/-^ shMcm6 CH12 cells**

The table contains the analysis of all genes analyzed using DESeq2^42^.

**Figure S1:**
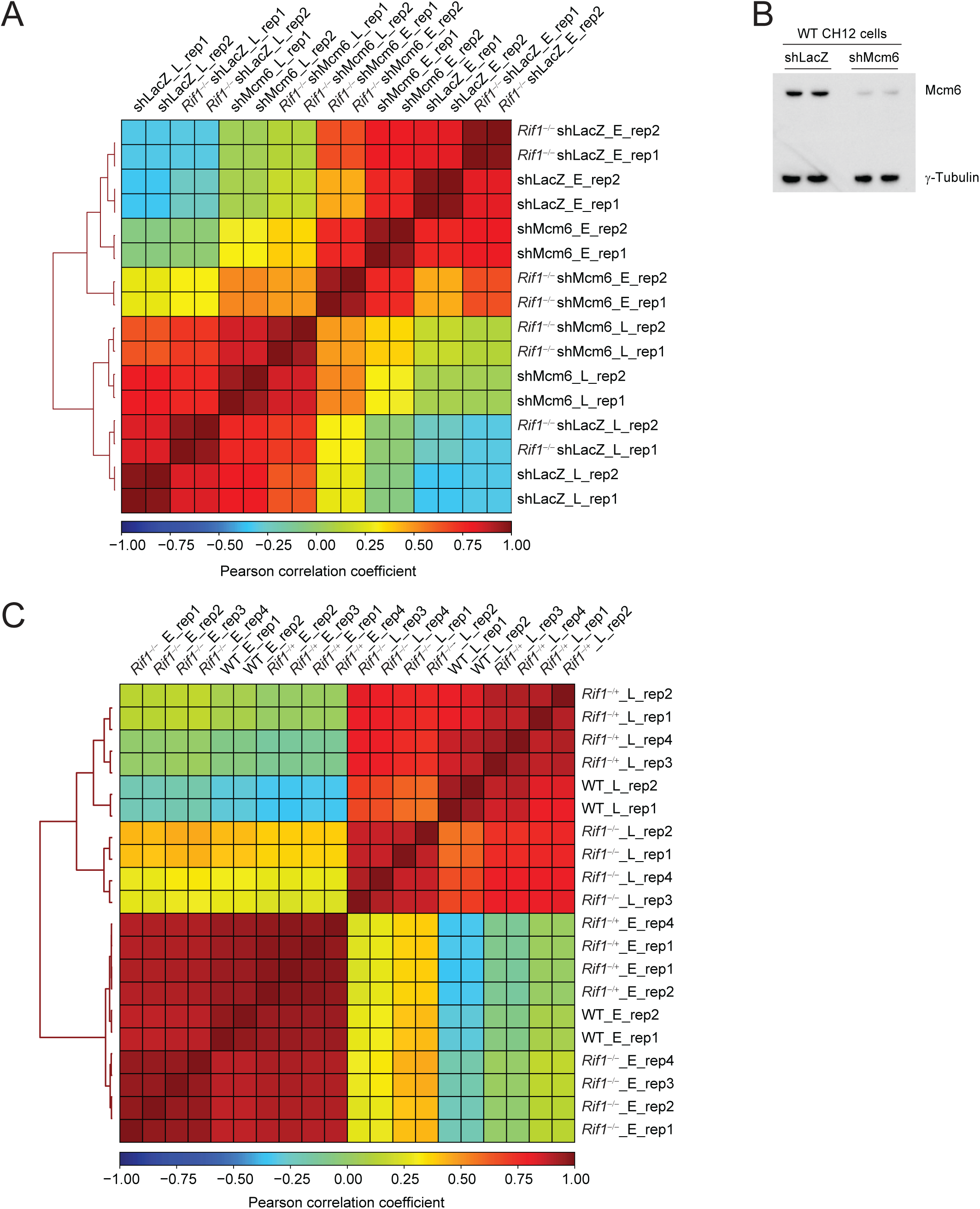
Analysis of Repli-seq replicates. A. Pearson correlation matrix of Repli-seq data from two replicates (rep1 and rep2) of shLacZ, shMcm6, *Rif1*^-/-^shLacZ and *Rif1*^-/-^ shMcm6 CH12 cells. E and L indicate the early and late fractions, respectively. The color gradient (bottom bar) corresponds to the Pearson correlation coefficient B. Western blot for Mcm6 protein levels from nuclear extracts obtained from shLacZ and shMcm6 cells. Data from two replicates are shown with γ-Tubulin serves as the loading control. In all cases, 20 μg of extract was loaded. C. Repli-seq correlation analysis as in A from WT (two replicates), *Rif1*^-/-^(four replicates) and *Rif1*^-/-^ (four replicates) primary, activated splenic B cells.

**Figure S2:**
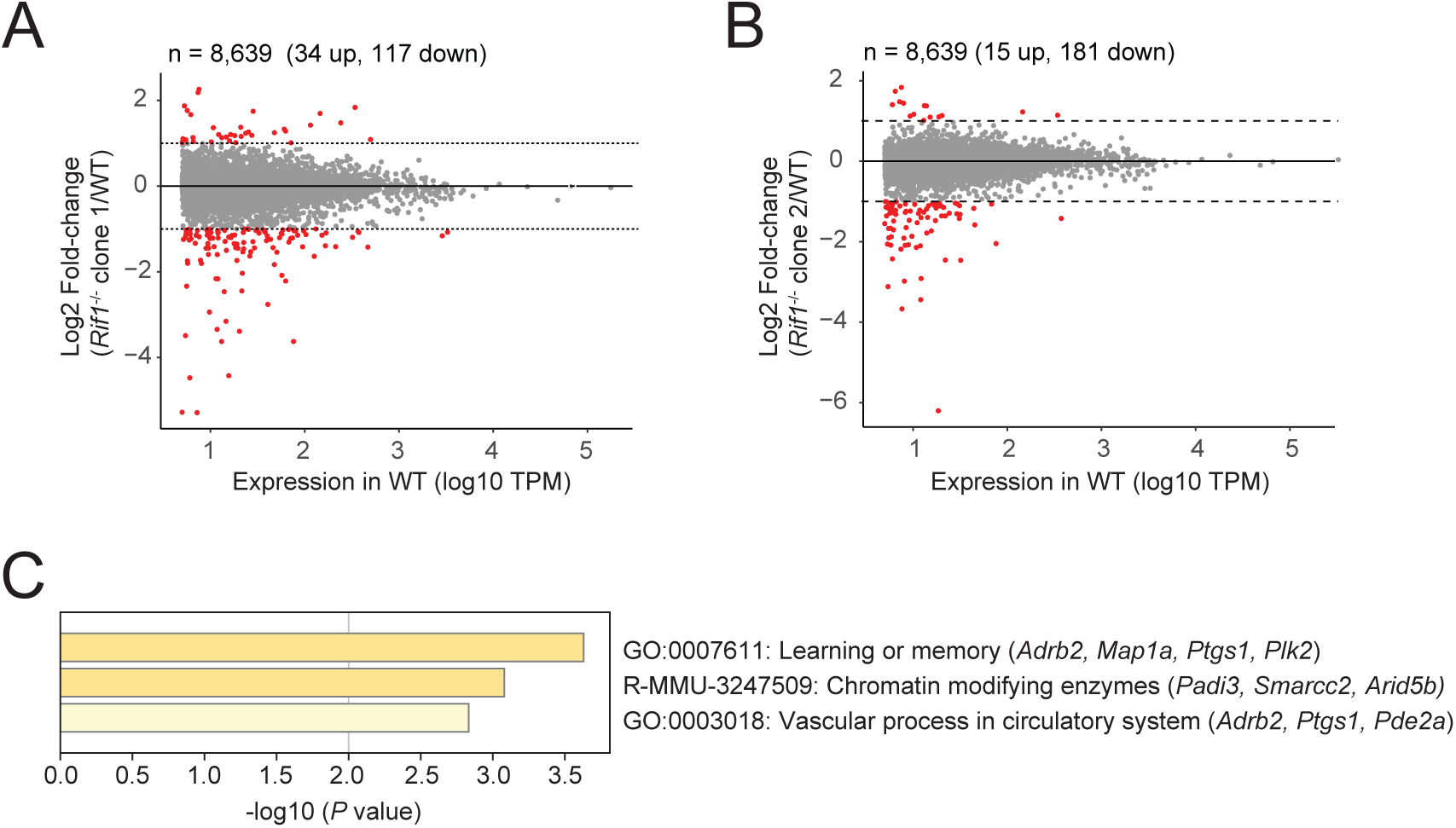
RNA-seq analysis from Rif1**^-/-^** CH12 cells. A. RNA-seq analysis of *Rif1*^-/-^ clone 1 compared to WT cells. The data are shown as a scatter plot of RNA-seq read densities in transcripts per million (TPM) versus fold-change (FC; *Rif1*^-/-^ /WT) for 8,639 expressed genes (defined as TPM > 5 in WT cells). The dotted lines mark log2 FC 1 or -1 corresponding to 2-fold upregulated and 2-fold downregulated genes, respectively. See also Table S1. B. As in A but for *Rif1*^-/-^ clone 2 compared to WT cells. C. GO term analysis of the downregulated genes common to *Rif1*^-/-^ clones 1 and 2 (n = 22). There were no enrichments for the common upregulated genes (n = 3). See also Table S1.

**Figure S3:**
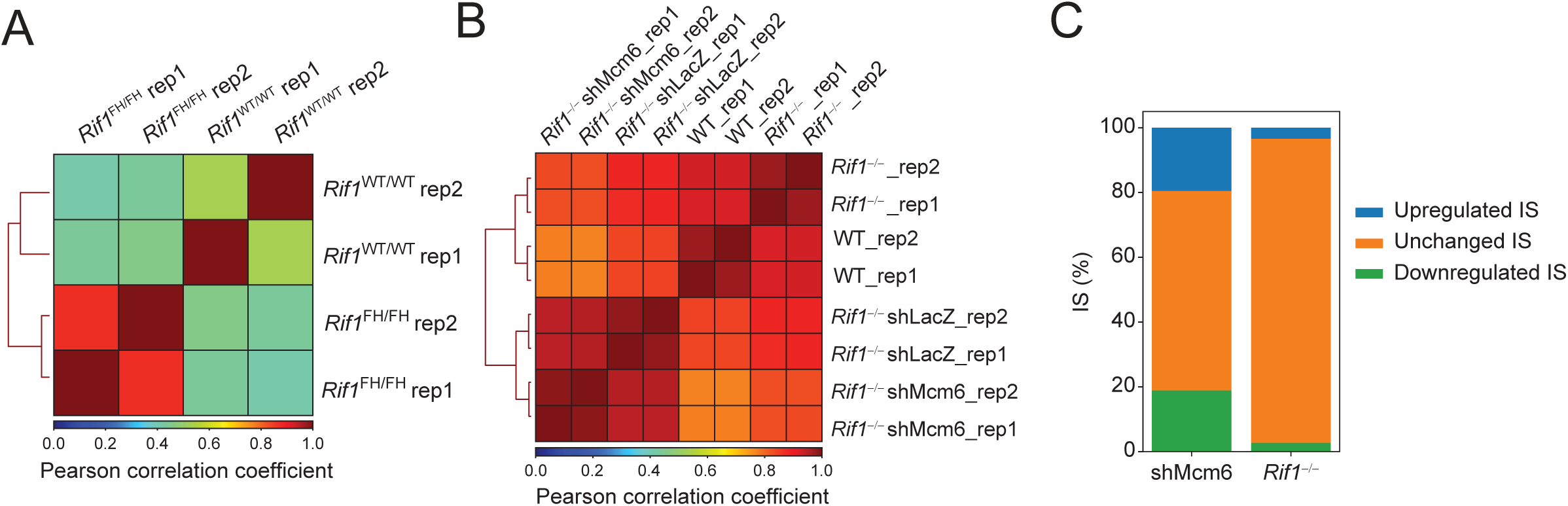
Analysis of ChIP-seq and SNS-seq replicates. A. Pearson correlation analysis of RIF1 ChIP-seq datasets. Two replicates from *Rif1*^FH/FH^ and *Rif1*^WT/WT^ primary B cells were analyzed. A. Pearson correlation analysis of SNS-seq data. Two replicates from each indicated condition were analyzed. B. Bar plots showing the percent of ISs upregulated (1.5-fold), downregulated (1.5-fold) or unchanged in shMcm6 (relative to shLacZ) or *Rif1*^-/-^ (relative to WT) cells. The shMcm6 data is from our previous study (ref. 13).

**Figure S4:**
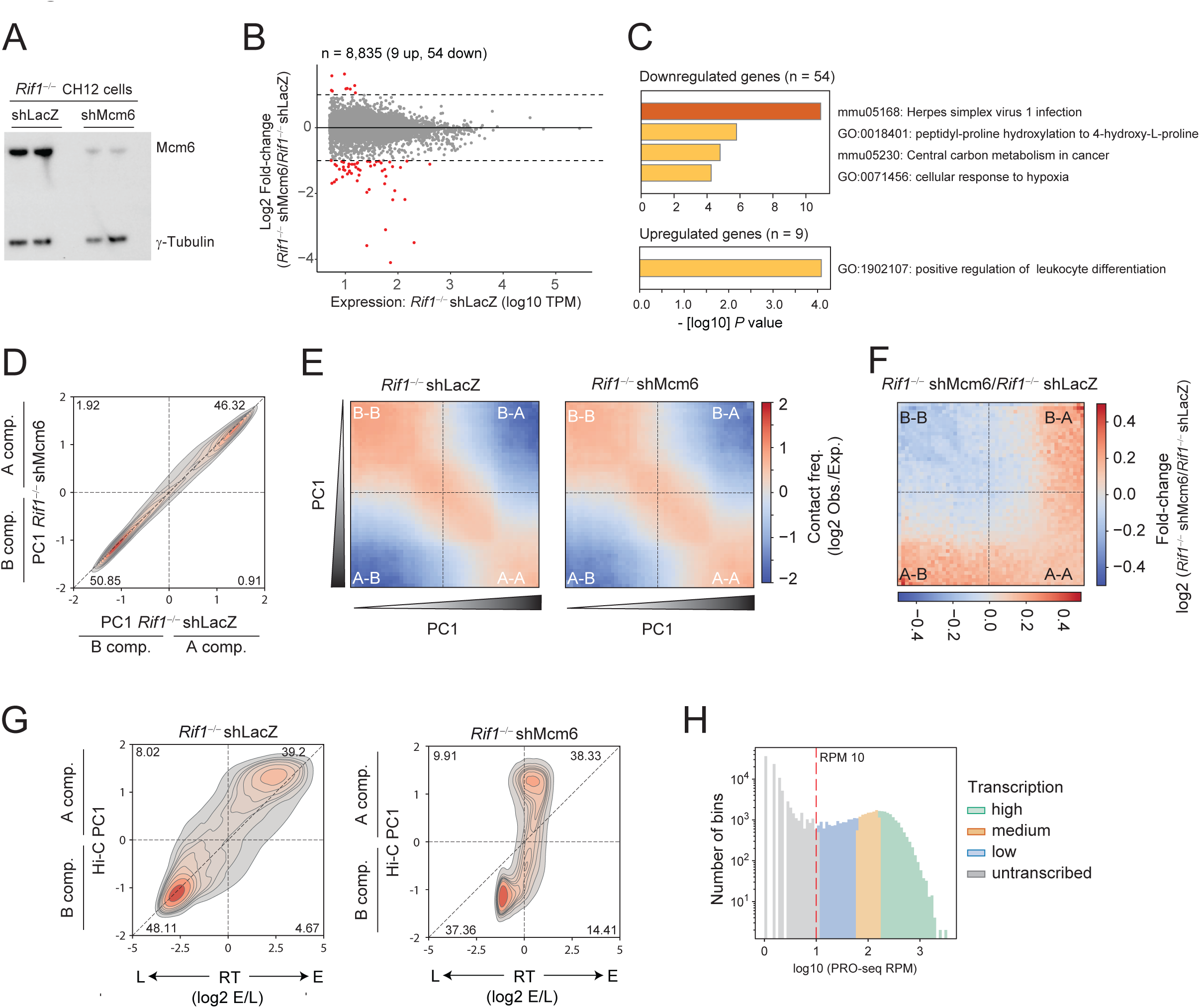
Gene expression and genome compartmentalization analyses in *Rif1***^-/-^** shMcm6 cells. A. Western blot for Mcm6 protein levels from nuclear extracts obtained from two replicates of *Rif1*^-/-^ shLacZ and *Rif1*^-/-^ shMcm6 cells. In all cases, 20 μg of extract was loaded. γ-Tubulin serves as the loading control. B. Scatter plot of RNA-seq read densities in TPM versus fold-change (*Rif1*^-/-^ shMcm6/Rif1*Rif1*^-/-^ shLacZ) at 8,835 expressed genes (defined as TPM > 5 in *Rif1*^-/-^ shLacZ cells). The dotted lines mark log2 FC 1 and -1 corresponding to 2-fold upregulated and 2-fold downregulated genes, respectively. C. GO term analysis of the 54 downregulated and 9 upregulated genes from B. D. Density-contour plot comparing PC1 compartment signals in 20 kb genomic bins from *Rif1*^-/-^ shLacZ and *Rif1*^-/-^ shMcm6 cells. E. Saddle plots comparing compartmental interactions between *Rif1*^-/-^ shLacZ and *Rif1*^-/-^ shMcm6 cells. F. Fold-change saddle plots based on data from E. G. PC1 versus RT density-contour plots in 20 kb genomic bins comparing the changes in either feature in *Rif1*^-/-^ shLacZ and *Rif1*^-/-^ shMcm6 cells. H. Histogram showing the distribution of PRO-seq densities (RPM) in 20 kb genomic bins from shLacZ cells. High, Medium, Low and Untranscribed groups are color-coded to match the analysis in Fig. 7B. The dotted red line indicates RPM 10 which marks the boundary between transcribed and untranscribed regions in our analyses.

